# The molecular principles governing HCMV infection outcome

**DOI:** 10.1101/2022.10.31.514490

**Authors:** Michal Schwartz, Miri Shnayder, Aharon Nachshon, Tamar Arazi, Yaarit Kitsberg, Roi Levi Samia, Michael Lavi, Rottem Kuint, Reuven Tsabari, Noam Stern-Ginossar

## Abstract

Infection with Human cytomegalovirus (HCMV) can result in either productive or non-productive infection, the latter potentially leading to establishment of latency, but the molecular factors that dictate these different infection outcomes are elusive. Macrophages are known targets of HCMV and considered to be permissive for productive infection, while monocytes, their precursors, are latently infected. Here we reveal that infection of macrophages is more complex than previously appreciated and can result in either productive or non-productive infection. By analyzing the progression of HCMV infection in monocytes and macrophages using single cell transcriptomics, we uncover that the level of viral gene expression, and specifically the expression of the major immediate early proteins, IE1 and IE2, is the principal barrier for establishing productive infection. On the cellular side, we reveal that the cell intrinsic levels of interferon stimulated genes (ISG), but not their induction, is a main determinant of infection outcome and that intrinsic ISG levels are downregulated with monocyte differentiation, partially explaining why macrophages are more susceptible to productive HCMV infection. We further show that, compared to monocytes, non-productive macrophages maintain higher levels of viral transcripts and are able to reactivate, raising the possibility that they may serve as latency reservoirs. Overall, by harnessing the tractable system of monocyte differentiation we decipher underlying principles that control HCMV infection outcome, and propose macrophages as a potential HCMV reservoir in tissues.

## Introduction

Human cytomegalovirus (HCMV) is a prevalent pathogen of the beta-herpesvirus family, infecting the majority of the human population worldwide ^1^. Although primary infection is usually asymptomatic in healthy individuals, HCMV can cause detrimental disease in immunocompromised individuals, and is the leading cause of congenital infections worldwide ^2^. Like all herpesviruses, HCMV can establish latent infection that persists for the lifetime of the host. Reactivation from latency in immunosuppressed individuals, such as transplant recipients and HIV patients, can result in severe morbidity and mortality.

HCMV infects a wide range of cell types including epithelial and endothelial cells, fibroblasts and myeloid cells ^2^. Infection outcome is tightly connected to cell type, and generally, differentiated cells are more permissive to productive infection, but the molecular determinants that govern HCMV permissiveness are still largely unknown. HCMV infection has been studied intensively in the myeloid lineage as cells along this lineage are thought to play a critical role in HCMV latency, reactivation, dissemination and persistence. CD34+ hematopoietic stem and progenitor cells (HSPCs), granulocyte-macrophage progenitor (GMPs) and blood monocytes are infected with HCMV but do not support productive infection and are the main cells in which HCMV latency has been characterized ^3–6^. Extensive work had been conducted on HCMV infected monocytes, illustrating infection activates cellular receptors and long term signaling events, promoting differentiation processes that support viral dissemination ^7, 8^. However, since CD14+ monocytes are short-lived, it has been proposed that the long term latent reservoir resides in bone marrow hematopoietic stem cells (HSCs) ^9^. It is noteworthy that the observed risk of HCMV reactivation in hematopoietic cell transplantations versus in solid organ transplantations ^10^ strongly implies that latent HCMV does not reside only in the bone marrow, and that there are additional uncharacterized sources of latent HCMV that likely reside within tissues ^11^.

Unlike monocytes, it was demonstrated that when macrophages and dendritic cells are infected with HCMV, they are permissive for productive infection ^12, 13^, and tissue macrophages were shown to be infected by HCMV *in vivo* ^14^. This difference in infection outcome between macrophages and their monocyte progenitors indicates a major role for the cellular environment, and repression of the virus via epigenetic mechanisms was shown to play an important role ^15^, however, it is not clear what drives these epigenetic differences and how the differences in infection outcome between different cell types is explained at the molecular level.

Furthermore, although macrophages are important for the first response to pathogens ^16^, reside in every tissue, the effect of HCMV infection in macrophages on their function is not well charted.

To dissect the molecular determinants that dictate infection outcome, we scrutinize HCMV infection of monocytes and macrophages at a single cell resolution. We reveal that macrophages can support both productive and non-productive infections and by comparing these two infection outcomes, as well as to infected monocytes, we demonstrate that the key determinant of infection outcome is the expression of viral immediate early genes. We show that non-productive infection is characterized by activation of Interferon-stimulated genes (ISGs), but this activation by itself does not deterministically affect infection outcome. On the other hand, we demonstrate that intrinsic ISG expression levels, independent of induction upon infection, are coupled to the differentiation state and have a substantial effect on infection outcome. We further reveal that productive infection perturbs macrophage cell identity and function. Finally, we show that non-productive macrophages are characterized by higher levels of viral gene expression compared to monocytes, and that the virus can reactivate from these cells. Altogether, our findings define determinants of HCMV infection outcome in the myeloid lineage, suggest that infection of macrophages interferes with their functions and raise the possibility that macrophages can serve as latent tissue reservoirs.

## Results

### Experimental system for studying HCMV infection outcome

In order to elucidate the molecular determinants that influence HCMV infection outcome, we sought to investigate the progression of infection in high temporal resolution in HCMV infected monocytes, which support latent infection, and in differentiated monocytes, which are permissive to productive infection. Initially, we infected CD14+ primary monocytes following different treatments that are used for differentiation, with HCMV TB40-E strain. The virus also contained a GFP reporter under the SV40 promoter ^17^, which allows quantification of productive infection efficiency. We tested three treatments that trigger primary monocyte differentiation in order to choose a strategy that will most efficiently support productive HCMV infection: phorbol-myristate (PMA) ^18–20^, granulocyte-macrophage colony-stimulating factor combined with interleukin-4 (GM-CSF/IL-4) ^20^ and macrophage colony-stimulating factor (M-CSF) ^21^. In addition, we tested TNFα treatment, which was previously shown to induce HCMV-permissive cells ^22^. As expected, 3 days post infection (dpi) of primary CD14+ monocytes with HCMV-GFP, almost all the cells show an increase in GFP indicating they are infected (Fig. 1a). However, GFP expression remains relatively low indicating that viral gene expression is repressed.

**Figure 1.**
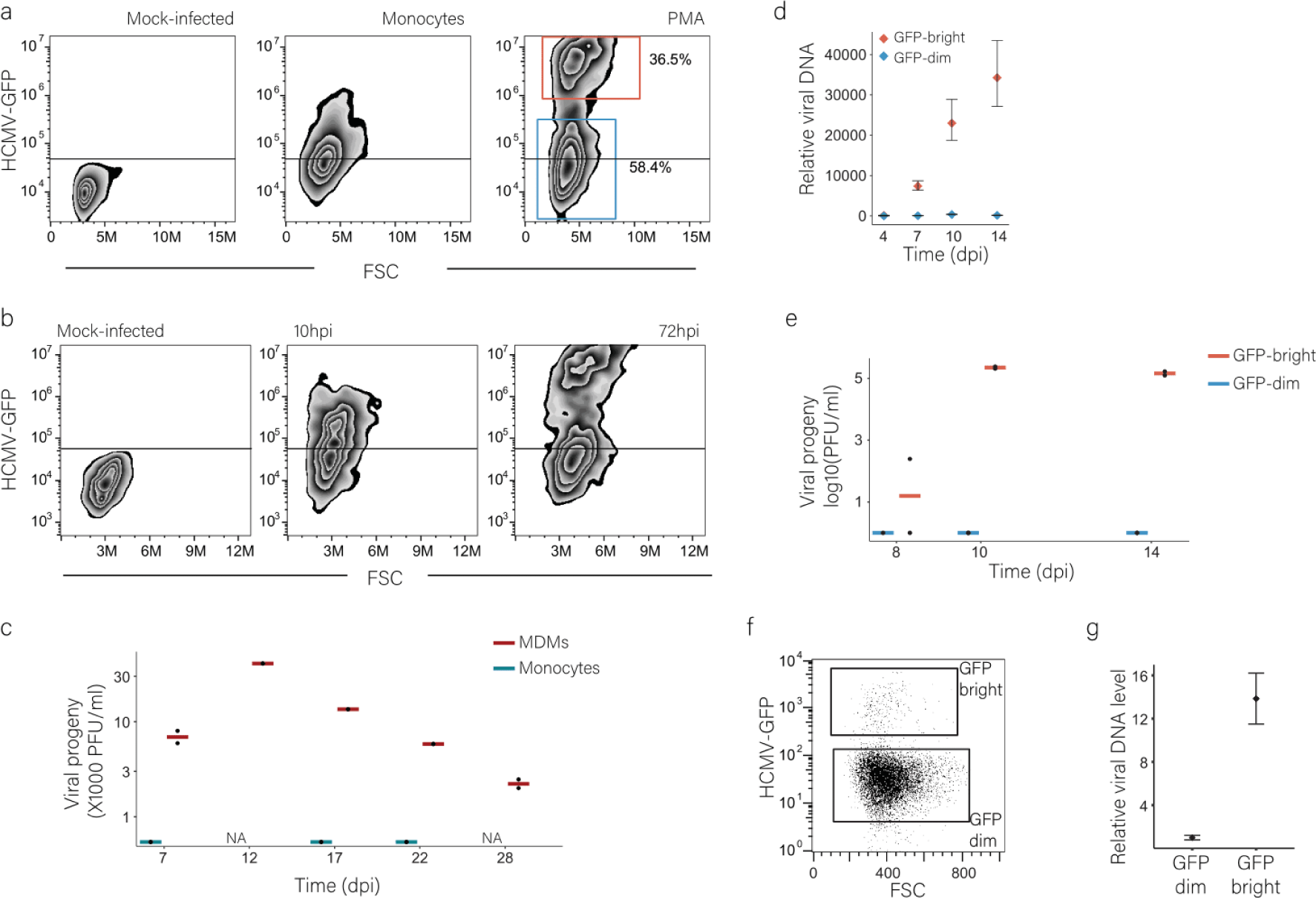
Productive and non-productive infection in monocyte derived macrophages. **a**. Flow cytometry analysis of primary CD14+ monocytes and monocytes treated with PMA, infected with HCMV-GFP. Analysis was performed at 3dpi. The mock-infected GFP level is marked by a line. Blue and red gates mark the GFP-dim and GFP-bright populations, respectively, and their percentage is noted. **b**. Flow cytometry analysis of PMA-induced monocyte-derived macrophages (MDMs) at different times following infection with HCMV-GFP. **c**. Measurements of infectious virus in supernatants from PMA-induced MDMs and from monocytes infected with HCMV-GFP at different times post infection. NA, not assayed. **d**. ddPCR measurements of viral genomes from FACS-sorted GFP-bright and GFP-dim HCMV-GFP infected MDMs at different times post infection. Graph is a representative of two biological replicates and shows the mean and 95% CV of poisson distribution. **e**. Measurements of infectious virus in supernatants from FACS-sorted GFP-bright and GFP-dim HCMV-GFP infected MDMs, at different times post infection. **f**. Flow cytometry analysis of BAL macrophages infected with HCMV-GFP. **g**. ddPCR measurements of viral genomes from FACS-sorted GFP-bright and GFP-dim HCMV-GFP infected BAL macrophages. Graph is a representative of two biological replicates and shows the mean and 95% CV of poisson distribution.

Interestingly, following all differentiation inducing-treatments, infection resulted in two distinct populations: one with a small increase in GFP levels (GFP-dim), suggesting these cells were infected with HCMV but like in monocytes, viral gene expression is repressed, and the second with a strong GFP signal (GFP-bright), suggesting productive infection (Fig. 1a and Extended Data Fig. 1a). Cells within each of the treatments displayed uniform shifts in surface expression of macrophage/dendritic cell markers, negating the possibility that differences in differentiation efficiency within the cell populations can solely explain the differences in infection outcomes (Extended Data Fig. 1b). For most further experiments we chose to use PMA derived macrophages as it is commonly used in HCMV studies ^19, 23–25^ and resulted in a large and distinct population of GFP-bright cells (Fig. 1a). Importantly, the two distinct populations in infected monocyte-derived macrophages (MDMs) are not apparent at early stages of infection, suggesting that they likely do not reflect two separate initial populations, but rather two different infection outcomes (Fig. 1b). Productive infection in MDMs was verified by detection of infectious virus in the supernatant, as long as several weeks post infection (Fig. 1c), despite ongoing cell death of cultured MDMs with time (Extended Data Fig 1c). In comparison, parallel infection of CD14+ monocytes yielded no infectious virus (Fig. 1c).

We sought to assess whether the two distinct GFP-expressing MDM populations seen by flow cytometry and microscopy indeed reflect different infection outcomes. First we noticed that GFP-bright cells are bigger compared to GFP-dim cells (Extended Data Fig. 1d and 1e), a known feature of lytic HCMV infection. We further sorted infected MDMs to GFP-bright and GFP-dim populations at 4 dpi (Extended Data Fig. 1f). Measurements of viral DNA levels show that at 4 dpi both the GFP-bright and the GFP-dim populations harbored viral DNA, yet viral DNA load was much higher in the GFP-bright cells suggesting viral DNA replication occurs in these cells (Extended Data Fig. 1g). Furthermore, viral DNA levels continued to increase with time in the GFP-bright population, while the GFP-dim population showed no substantial change (Fig. 1d). These results indicate that viral genome replication was taking place in the GFP-bright population but not in the GFP-dim. In addition, supernatants from GFP-dim and GFP-bright MDMs were collected at different dpi and used to infect fibroblasts in order to measure production of infectious virus. We failed to detect any infectious virus in the supernatant of the GFP-dim cells at any time point. However, measurable infectious virus was detected in the supernatant of the GFP-bright cells, reaching highest titers at 10dpi (Fig. 1e). These results indicate that the majority of MDMs were infected with the virus but a productive infection only developed in part of the population. The split of infected macrophages into two infection outcomes occurs also in MDMs differentiated with all other treatments tested, as we could verify the discernible GFP bright and dim populations differed in viral genome levels at 4 dpi (Extended Data Fig. 1 h-j). To explore whether these two infection outcomes occur also in natural tissue myeloid cells, we collected cells from BronchoAlveolar Lavage (BAL) samples and infected them with HCMV-GFP. The presence of macrophages in these samples was verified by size, morphology and surface marker staining (Extended Data Fig. 1k and 1l). Indeed also in BAL macrophages we observed two distinct populations of GFP-dim and GFP-bright cells that differ in viral genome levels (Fig. 1f, 1g and Extended Data Fig. 1l).

### Single-cell transcriptome analysis along HCMV infection of CD14+ monocytes and MDMs

In order to shed light on the viral and cellular processes governing these two infection outcomes, we turned to single-cell RNA-sequencing (scRNA-seq). Differentiation of primary monocytes provided a tractable system which allowed a head to head comparison between infection in monocytes and macrophages and a large enough number of cells to be able to analyze infection at multiple time points. We analyzed infected primary CD14+ monocytes and PMA-induced MDMs from the same donor at 5, 12, 24, 48, 72 and 144 hours post infection (hpi), as well as mock-infected CD14+ monocytes and MDMs at the 5 and 144 hpi time points (Fig. 2a). In total we analyzed 871 monocytes and 2355 MDMs. Projection of all cells according to both cellular and viral reads, revealed several distinct groups of cells (Fig. 2b and 2c).

**Figure 2.**
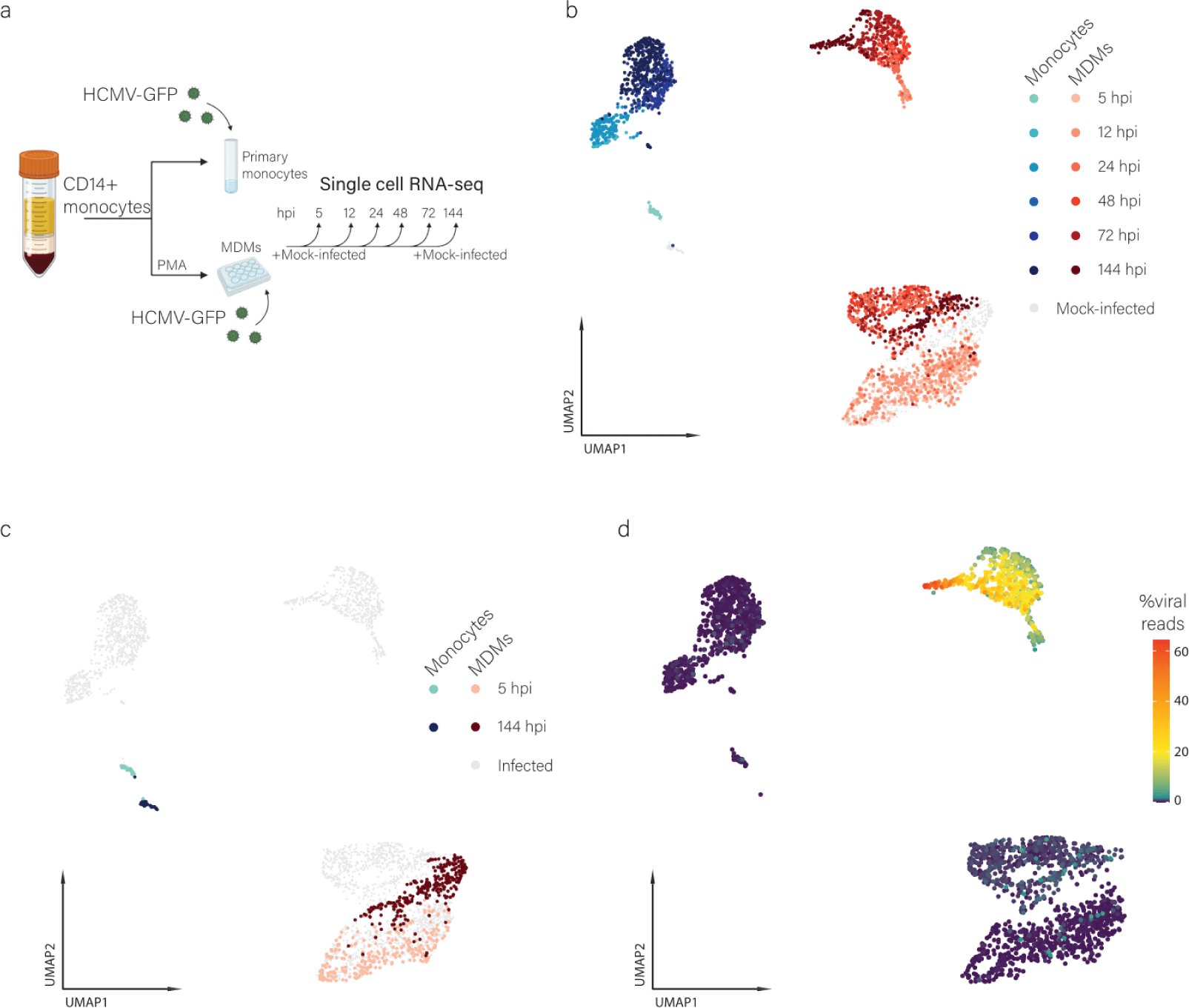
Single-cell transcriptome analysis in high temporal resolution of HCMV infection of primary monocytes and MDMs. **a**. Experiment design. primary monocytes and PMA-induced MDMs were infected with HCMV- GFP and at 5, 12, 24, 48,72 and 144 hpi were processed for scRNA-seq. In parallel mock-infected cells from 5 and 144 hpi were also processed for scRNA-seq. **b and c**. Projections of HCMV-GFP infected CD14+ monocytes (n=803) and MDMs (n=1815) as well as mock-infected monocytes (n=68) and MDMs (n=540) from the indicated time points, based on cellular and viral gene expression, colors indicate cell types and time post infection. The colored cells show either the infected cells (**b**) or the mock-infected cells (**c**). **d**. Projection as in b, showing all infected cells, colored by percentage of viral reads per cell.

At 5 hpi, infected monocytes were grouped with mock-infected monocytes, indicating that transcriptional changes at this time point are milder compared to later times in infection (Fig. 2b and 2c). Infected monocytes from 12 hpi onwards appeared as a single population (Fig.2b) and the cellular genes that dictated their stratification from early monocytes were mainly interferon stimulated genes (Extended Data Fig. 2). As expected, viral transcript load in infected monocytes remained low along infection (median=0.063%), and did not exceed 1% of the total transcriptome (Fig. 2d), indicating lack of productive infection, in line with previous reports ^26–28^. In contrast to monocytes, and in agreement with our flow cytometry analysis, infected MDMs from 24 hpi and onwards, were separated into two distinct populations (Fig. 2b) that were characterized by drastic differences in viral transcript load (Fig. 2d). These two populations reflect two infection outcomes: in one, viral transcript load increased as infection progressed, reaching more than 50% of the transcriptome, illustrating these cells represent productive infection. The second group of MDMs was composed of mock-infected MDMs, most of the infected MDMs from the early time points, as well as 57% of MDMs at the later time points. In infected MDMs that belong to this group, viral transcript loads were low (median=0.226%) and did not dramatically differ from infected monocytes (Fig. 2d).

### Two distinct infection outcomes in infected macrophages

We next focused on analyzing infected MDMs separately, as the two separate populations give an opportunity to explore the trajectories that lead to these two distinct infection outcomes. Notably, all mock-infected MDMs were transcriptionally distinct from the initial monocyte population (Extended Data Fig. 3a) and showed an increase in macrophage differentiation signature (Extended Data Fig. 3b) indicating that it is unlikely that the two distinct infection outcomes originate from two initially distinct populations. Projection of all infected MDMs according to both cellular and viral transcripts, revealed that cells were divided into three discernible populations, stratified further into sub-clusters (Fig. 3a). Most of the cells from the early time points (5-12 hpi) belong to a single group, while cells from later time points in infection split into two additional distinct groups (Fig. 3b), which differ greatly in the level of viral transcripts (Fig. 3c). These divisions illustrate that initially, all cells start with low levels of viral transcripts and are indistinguishable in regard to infection outcome. Beginning at 12, and in most cases at 24 hpi, cells transition to one of two groups: either productive infection, as indicated by the high levels of viral transcripts, reaching 20%-70% at late times of infection, or non-productive infection in which viral transcript levels remain low (Fig 3d and Fig. 3e). These features,therefore, define three main populations of infected MDMS: early, non-productive and productive (Fig 3d). Clustering further stratified cells in each of these populations (Fig 3a), which corresponded to progression in the given trajectory with time (Fig. 3b and 3d). The productive and non-productive populations, showing low and high viral loads, correspond to low and high GFP expression (Extended Data Fig. 3c), indicating that the two distinct GFP populations indeed reflect the two different infection outcomes in MDMs.

**Figure 3.**
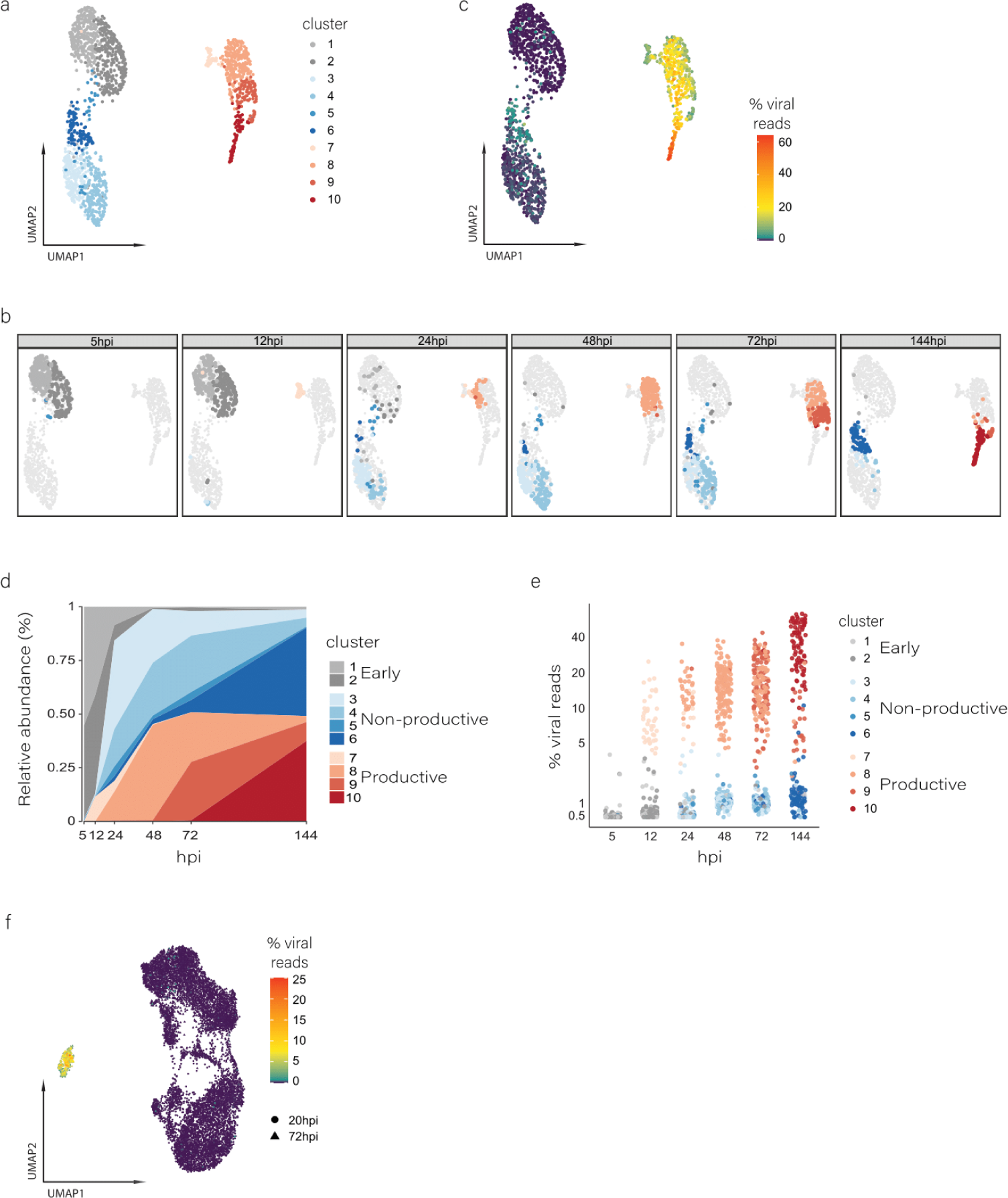
Single-cell transcriptome analysis of HCMV infected macrophages. **a**. Projection of 1815 HCMV-GFP infected MDMs at 5-144 hpi colored by cluster assignment. Analysis was based on both cellular and viral transcripts. **b**. Projections as in figure 3a, showing the cells from each time point along infection. **c**. Projection, as in figure 3a, colored by percentage of viral reads per cell. **d**. Distribution of the single cells between the different early, non-productive and productive clusters along infection. **e**. Percentage of viral reads in single MDMs from the different clusters along infection. **f**. Projection of 10,320 HCMV-GFP infected myeloid BAL cells from 20 and 72 hpi colored by percentage of viral reads per cell. Circles and triangles mark cells from 20 or 72 hpi, respectively. Analysis was based on both cellular and viral transcripts.

We further applied scRNA-seq to total infected BAL cells. Since the number of cells was very low, we sampled the cells only at two time points post infection: 20 hpi and 72hpi. Projection of the cells according to both cellular and viral transcripts, demonstrated that the cells from both time points stratify into two distinct populations of cells and an additional small population that expresses significantly higher viral transcript levels and reflects productively infected cells (Extended Data Fig. 3d and 3e). Analysis of marker genes showed one population exhibits signatures of cytotoxic lymphocytes, with genes encoding granzymes and T/NK cell related surface markers and cytokines, whereas the second population represented myeloid cells/macrophages, with genes related to antigen presentation and phagocytosis (Extended Data Fig. 3f). The productive cells also expressed myeloid cell markers (Extended Data Fig. 3f), demonstrating these cells originate solely from the myeloid cells. Therefore, in order to focus on the relevant signatures to HCMV infection outcome, we focused only on the myeloid cells. As in the MDMs, we observe a distinct group of productively infected cells, as evident from the much higher levels of viral transcripts and another group, in which the cells carry low levels of viral transcripts (Fig. 3f).

### Expression levels of viral immediate early genes govern infection outcome

In order to delineate the contribution of cellular and viral genes to the stratification of the MDMs to the different groups (early, non-productive, productive), we next analyzed the transcriptome solely according to cellular genes. This analysis produced the same three distinct groups of cells and very similar cluster stratification, indicating these three infection states are associated with corresponding changes in the cellular transcriptome (Fig. 4a and Extended Data Fig. 4a). The exceptions were clusters 5 and 7. Cluster 5 includes very few non-productive cells from different time points and these cells clustered with early cells when clustering was based only on cellular genes. The nature of these cells is unclear but since this cluster is very small these cells were removed from following cluster-based analyses.

**Figure 4.**
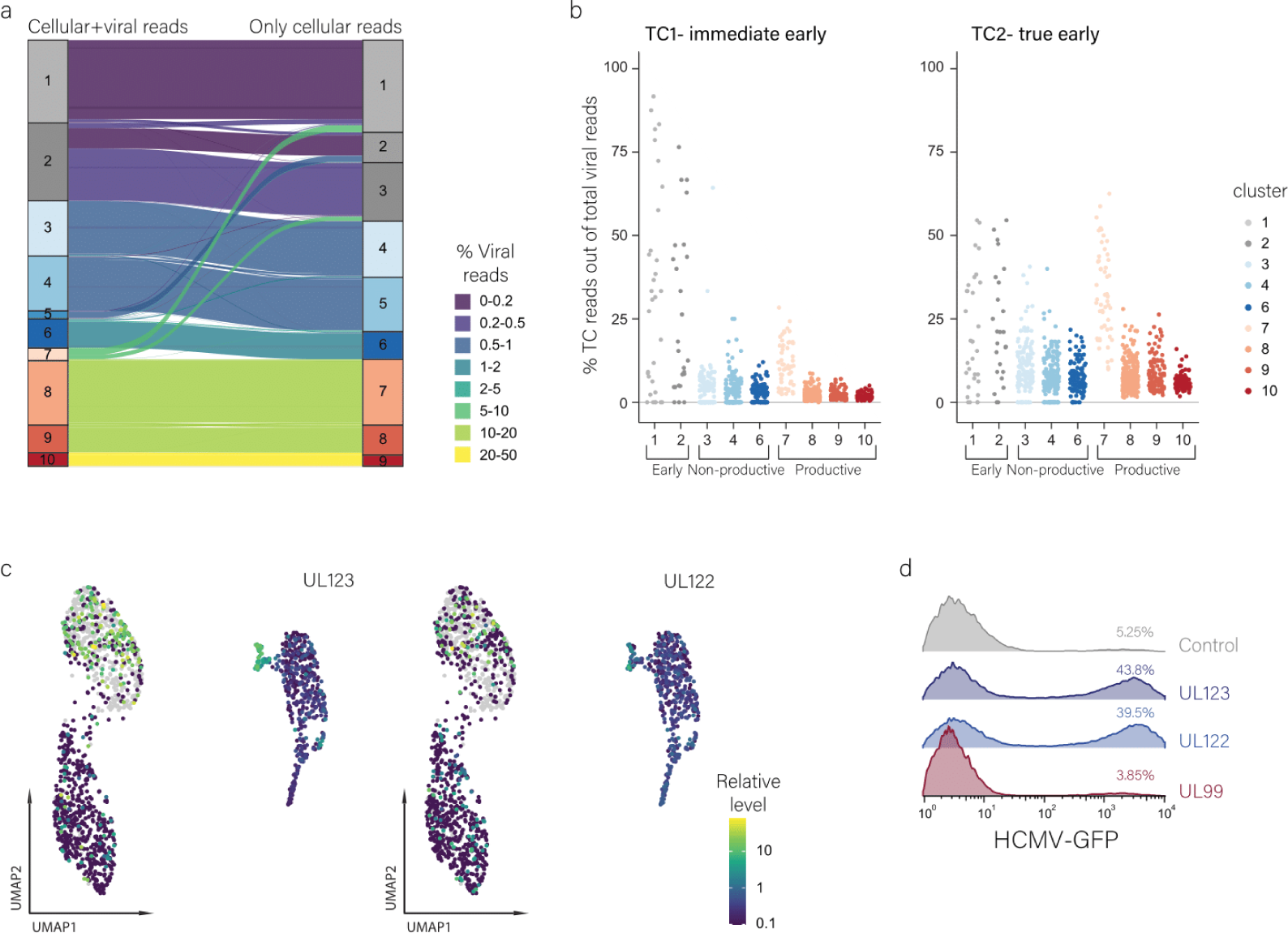
Expression of viral immediate early genes govern infection outcome of MDMs. **a**. Cluster assignment of each cell according to analysis based on both viral and cellular genes (left) or on analysis based solely on cellular genes (right). Each cell, represented by a single line, is colored by the percentage of viral reads. **b**. Relative expression levels, in the different clusters, of viral genes from temporal classes 1 (immediate early genes) and 2 (true early) based on ^28^. **c**. Projection of infected MDMs (as in Fig. 3a) colored by the log expression level of UL123 or UL122 relative to all other viral genes. **d**. Flow cytometry analysis of HCMV-GFP infected THP1- derived macrophages over expressing mCherry-flag as control, UL123, UL122 or UL99. Replicate data and statistics are shown in Extended Data Fig. 4e.

Cluster 7 is the earliest productive cluster and contains cells that already express substantial levels of viral transcripts (median= 8.7%, Fig. 3a). When clustering was based only on cellular genes, these cells clustered together with the early cells, showing that their classification as productively infected is dictated mainly by viral gene expression (Fig. 4a). Also when infected BAL myeloid cells were analyzed based only on cellular genes it became clear that some of the cells that are productively infected are defined as such only based on the expression of viral genes (Extended Data Fig. 4b). This demonstrates that the major first transcriptional signature that defines cells as productively infected is robust viral gene expression. We next examined which viral genes are expressed at the transition from early to productive cells. For this, we took advantage of the recent temporal classification of HCMV gene expression ^28^. We found that the distinction of these early productive cells (cluster 7) was the high relative expression of IE genes (TC1, Fig. 4b), as well as specific induction of true early genes (TC2, Fig. 4b), compared to the other viral temporal classes (Extended Data Fig. 4c). This suggests that expression of IE genes at a level which is sufficient to initiate the induction of early genes, is the main determinant that dictates the progression towards productive infection. Examination of the relative expression levels of the two transcripts transcribed from the major immediate early promoter, UL123, which encodes IE1, and UL122, which encodes IE2, showed that they indeed have specific expression in these early productive cells (Fig. 4c). To test if the initial expression levels of UL123 or UL122 indeed dictate HCMV infection outcome, we ectopically expressed either mCherry (that served as a control), UL123, UL122 or UL99, a viral late protein that was used as an additional control, in macrophages derived from the monocytic cell line THP1. To avoid prolonged expression of the viral proteins, we used a doxycycline-inducible system and induced the expression of the transgene only 24 hour prior to infection, after the cells were differentiated. Similar to infection with primary monocyte-derived MDMs, infection of THP1 MDMs resulted in two populations of GFP-bright and GFP-dim cells, representing productive and non-productive infection, respectively (Extended Data Fig. 4d). In accordance with the signature revealed by our single cell analysis, we show that ectopic expression of either UL123 or UL122 but not UL99 significantly increases the percentage of productively infected cells (Fig. 4d and Extended Data Fig. 4e), demonstrating that the expression of IE genes is indeed a major barrier that dictates infection outcome in MDMs. Notably, following overexpression, levels of the corresponding viral proteins were substantially increased (Extended Data Fig. 4f and 4g). The effect of ectopic expression of UL123 was further validated in another commonly used model for HCMV infection, PMA treated Kasumi-3 (Extended Fig. 4h).

Since scRNA-seq gives unprecedented detail on inter-cellular heterogeneity in gene expression profiles, we next explored whether we can use this heterogeneity to identify cellular gene expression profiles that are associated with the initial levels of viral gene expression . Calculation of the correlation coefficient between each cellular transcript and either viral gene expression or specifically with UL123 and UL122 expression, in early infected MDMs, did not reveal any cellular genes whose expression significantly correlates or inverse correlates with the initial levels of viral genes (Extended Data Fig. 4i). Our inability to detect variations that are associated with induction of viral gene expression, likely reflects the relatively homogenous nature of the cells we used and therefore small differences in gene expression are likely masked by other factors, such as the amount of incoming virions which varies between cells ^29^, or variability between the content of virions ^30^, both which likely play a major role in setting infection outcome.

### Induction of interferon stimulated genes does not dictate infection outcome

We next examined the features that characterize the shift from early infection to the non-productive population. The cellular genes that were most upregulated in the non-productive cells were Interferon-Stimulated Genes (ISGs, Extended Data Fig. 5a). These genes were strongly upregulated in the non-productive cells and barely in the productive cells (Fig. 5a and 5b). Since the main characteristics of non-productive and productive cells are the induction of ISGs and induction of viral gene expression, respectively, we next examined the relationship between these two features in single cells along MDM infection. This analysis revealed that at early time points, both viral gene and ISG expression are low. As time progressed, most of the cells increased either viral gene expression or ISG expression (Fig. 5c). Notably, the cells of both trajectories were exposed to the same extracellular signals (e.g. interferons), as they were grown together in the same culture dish. This indicates that the induction of viral gene expression, and likely the specific induction of IE genes and/or additional early genes in the productive cells, renders the cells unresponsive to IFN signaling. Since we hardly detected infected MDMs that induced both viral genes and ISGs, this analysis may imply that these events describe two competing deterministic outcomes - induction of viral genes resulting in productive infection, or induction of ISGs prohibiting productive infection. However, analysis of the percentage of cells in the different populations at each time point indicates that productive cells must also originate from the non-productive groups in which ISGs were already induced, as the percentage of productive cells reaches numbers that can not be solely explained by transition from the early infected population (Fig. 5d). This therefore suggests that induction of a robust interferon response does not forestall establishment of productive infection and the main determinant that governs infection outcome is the induction of viral gene expression above a certain threshold. Analysis of a recent single cell RNA-seq dataset along HCMV infection of fibroblasts ^31^ demonstrated the same phenomenon. At 96 hpi, a time point that permits mostly a single round of infection, almost all fibroblasts are productively infected, nevertheless a significant portion of the cells went through a stage in which ISG expression was induced and viral gene expression was low, at earlier time points (Extended Data Fig. 5b). These observations in two independent systems suggest that induction of ISGs by an infected cell does not deterministically prevent the establishment of productive infection within that cell.

**Figure 5.**
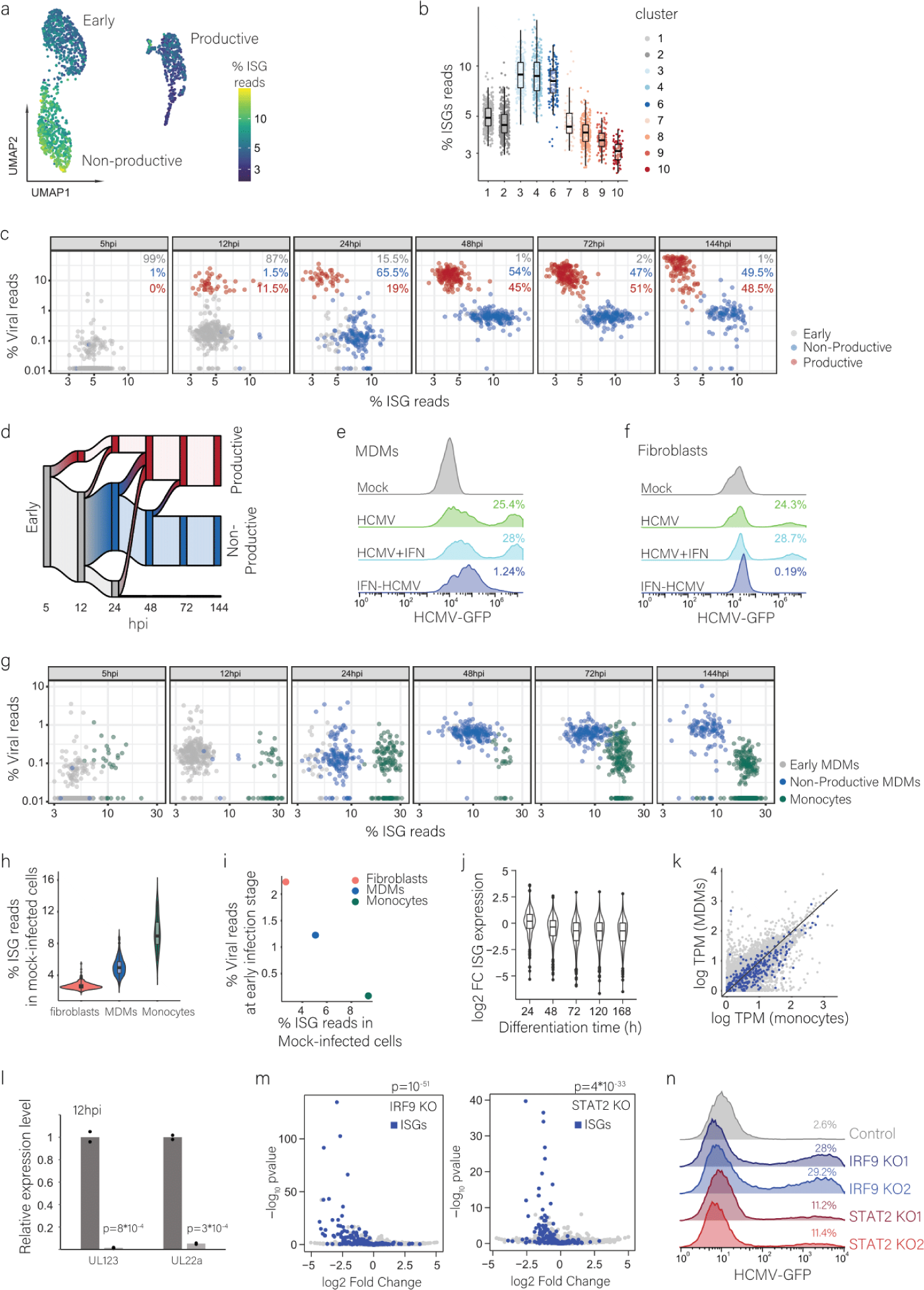
Expression of ISGs prior to infection but not their induction following infection affects infection outcome. **a**. Projection of infected MDMs (as in Fig. 3a) colored by the expression level of ISGs. **b**. Expression level of ISGs in single MDMs from the different clusters. **c**. Viral gene expression level versus ISG expression level, in single early, non-productive and productive MDMs at different times post infection. The percentage of each group of cells in the given time point is indicated. **d**. Flow chart describing the transition of cells between early, non-productive and productive groups along infection. **e**. and **f**. Flow cytometry analysis of HCMV-GFP infected MDMs (**e**) or fibroblasts (**f**) either left untreated, treated with interferons (IFNs) at the time of infection (HCMV+IFN) or treated with IFNs 24 hours prior to infection (IFN-HCMV). Mock-infected cells are shown as control. Replicate data and statistics are shown in Extended Data Fig. 5c, 5e, 5g and 5h. **g**. Viral gene expression level versus ISG expression level in single infected monocytes, early and non-productive MDMs at different times post infection. **h**. ISG expression level in mock-infected monocytes and MDMs as measured from the scRNA-seq data and in mock- infected fibroblasts based on data from ^31^. **i**. ISG expression level in mock-infected cells versus viral gene expression level at early stages of infection (12 hpi or 20 hpi, in monocytes/MDMs and fibroblasts, respectively). Data is based on the current sc-RNA-seq data for monocytes and MDMs and on ^31^ for fibroblasts. **j**. Expression levels of different ISGs in cells along differentiation trajectory from monocytes to MDMs. Data taken from ^32^. **k.** Cellular transcript levels in monocytes and MDMs. Data taken from ^32^. Blue dots mark ISGs. TPM, transcript reads per kilobase per million reads mapped to cellular transcripts. **l.** Relative expression level of the viral transcripts UL123 and UL22A, measured by qPCR, in infected MDMs either untreated or treated with interferons (IFNs) 24 hours prior to infection, at 12hpi. Points show measurement of biological replicates. p-values as calculated by t-test on the replicates are indicated. **m.** Differential expression analysis of cellular genes in uninfected THP1 derived macrophages (DMs) following CRISPR KO of IRF9 (left panel) or STAT2 (right panel) compared to control CRISPR KO. Blue dots mark ISGs. **n.** Flow cytometry analysis of HCMV-GFP infected THP1 DMs following CRIPSR mediated knockout of a control gene, IRF9 or STAT2. Two independent gRNAs were used for IRF9 and STAT2. Percentage of productively infected cells are indicated for each condition. Replicate data and statistics are shown in Extended Date Fig. 5n.

To experimentally examine whether IFN response has deterministic effects on infection outcome, we tested whether boosting the IFN response co-currently with infection will affect infection outcome in MDMs. In accordance with the results of our analysis, adding IFNs (500U/ml IFNα, 700U/ml IFNβ and 500U/ml IFNγ) at the time of infection did not affect the percentage of productively infected cells (Fig. 5e and Extended Data Fig. 5c), despite significant increase in ISG expression (Extended Data Fig 5d). Similar results were obtained in low MOI infection of fibroblasts, which are bona fide lytic infection supporting cells (Fig. 5f and Extended Data Fig. 5e). Together these results demonstrate that activation of ISGs in response to infection may slow productive infection but overall does not deterministically prevent it.

### Intrinsic ISG levels affect HCMV infection outcome

We next examined viral gene and ISG expression in monocytes. As expected, all monocytes exhibit low viral gene expression levels which is comparable at earlier stages to infected MDMs (Fig. 5g). Notably, compared to MDMs, the expression level of ISGs was substantially higher in monocytes even at the earliest stage of infection (Fig. 5g). The substantial differences in ISG levels between monocytes and MDMs at the very early time point of infection, which likely proceed the induction of IFN, prompted us to examine if there are differences in the intrinsic levels of ISGs and whether they relate to the permissiveness to productive HCMV infection. We first examined the scRNA-seq data of mock-infected monocytes and MDMs from this study, as well as fibroblasts from Hein et al. ^31^. We found that indeed the intrinsic levels of ISGs (in mock-infected cells) in fibroblasts is lower than in MDMs, which in turn is lower than in monocytes (Fig. 5h) and there is a clear inverse-correlation between the intrinsic levels of ISGs in mock-infected cells and the levels of viral gene expression at early time points of HCMV infection (Fig. 5i). To examine this finding along differentiation of monocytes, we analyzed published transcriptomic data along the differentiation trajectory from monocytes to MDMs ^32^ which revealed a decrease in the intrinsic levels of many ISGs along this trajectory (Fig. 5j and Fig. 5k). Since the permissiveness for lytic infection is tightly linked to differentiation state ^33, 34^, we examined the link between intrinsic expression of ISGs and differentiation state of hematopoietic cells in the bone marrow. Analysis of two scRNA-seq datasets ^35, 36^ revealed that intrinsic ISG levels are reduced as cells differentiate (Extended Data Fig. 5f). These results suggest a tight link between basal levels of ISGs, differentiation and permissiveness to productive HCMV infection.

To explore if the levels of ISGs at the time of infection affect infection outcome, we tested the effect of IFN pre-treatment on MDMs and fibroblasts on HCMV infection. Indeed, addition of IFNs 24 hours prior to infection almost completely eliminates productive infection in both MDMs and fibroblasts (Fig. 5e, Fig. 5f, extended Fig. 5g and 5h). IFN pretreated cells did not produce infectious progeny up to 10 days post infection (Extended Data Fig. 5i) and viral DNA replication was completely blocked (Extended Data Fig. 5j) indicating a blockage of productive infection and not a mere delay. We further show that in IFN pretreated cells viral gene expression is extremely reduced already at 12hpi (Fig. 5l), indicating that ISGs act to block the virus at the stage of immediate early viral gene expression or earlier.

While these experiments show that boosting ISG expression prior to infection affects infection outcome, we next sought to test if intrinsic ISG levels, without any boost, also play a role. Intrinsic levels of ISGs may originate from low basal levels of interferon signaling, but it was recently suggested that basal expression of ISGs is controlled by preformed STAT2-IRF9 complexes, which are independent of interferon receptors signaling ^37^. To test these possibilities we first treated MDMs, 24 hours prior to infection, with ruxolitinib, a potent and selective Janus kinase (JAK) 1 and 2 inhibitor ^38^ that blocks the signaling downstream of interferon receptors.

This treatment did not affect the intrinsic levels of ISG (Extended Data Fig. 5k) and correspondingly did not affect infection outcome (Extended Data Fig. 5l). We therefore next tested whether we could reduce intrinsic ISG levels by depleting IRF9 or STAT2 using CRISPR- cas9 in THP1 derived macrophages (Extended Data Fig. 5m). In support of their role in regulating basal ISG expression independently of IFN, depleting either IRF9 or STAT2 reduced the basal expression of many ISGs (Fig. 5m). Importantly, depletion of IRF9 or STAT2 using two independent guide-RNAs increased the percentage of productively infected cells (Fig. 5n and Extended Data Fig. 5n). Furthermore this effect was not diminished by addition of ruxolitinib at the time of infection (Extended Data Fig. 5o), illustrating the increase in productive cells is due to the reduction in intrinsic ISGs rather than reduced induction of ISGs following infection.

Overall these results point to a major role for intrinsic ISG expression in dictating HCMV infection outcome and suggest that the intrinsic expression of ISGs, that decrease upon differentiation, contributes to the permissiveness of differentiated cells to HCMV infection. Furthermore, by combining differential ISG expression in our various experiments, we highlight ISGs that may be more likely to be involved in determining HCMV infection outcome (Supplementary Table 1) and future work is needed to delineate their specific effects.

### Productive HCMV infection disrupts macrophage identity and function

To characterize cellular transcriptional changes following the different infection outcomes in MDMs, we performed differential gene expression analysis comparing MDMs from the different infection trajectories to mock-infected cells, followed by pathway enrichment analysis. Several metabolism pathways that are known to be enhanced during HCMV infection in fibroblasts ^39^ were significantly enriched in productive MDMs, including the TCA cycle and fatty acid metabolism (Fig. 6a). In line with the virus subverting the cell cycle machinery to its own benefit ^40^, several pathways related to modulation of the cell cycle were enriched. In addition, HCMV is reliant on the cellular machinery for expression of its genes, as reflected by modulation of different pathways related to gene expression with a strong induction of genes related to translation ^41, 42^. As mentioned above, non-productive MDMs significantly induced, mainly and most robustly, ISGs. These results demonstrate that the non-productive trajectory is dominated by induction of ISGs, while in the productive trajectory the virus takes over and subverts the cellular machinery for promoting its propagation.

**Figure 6.**
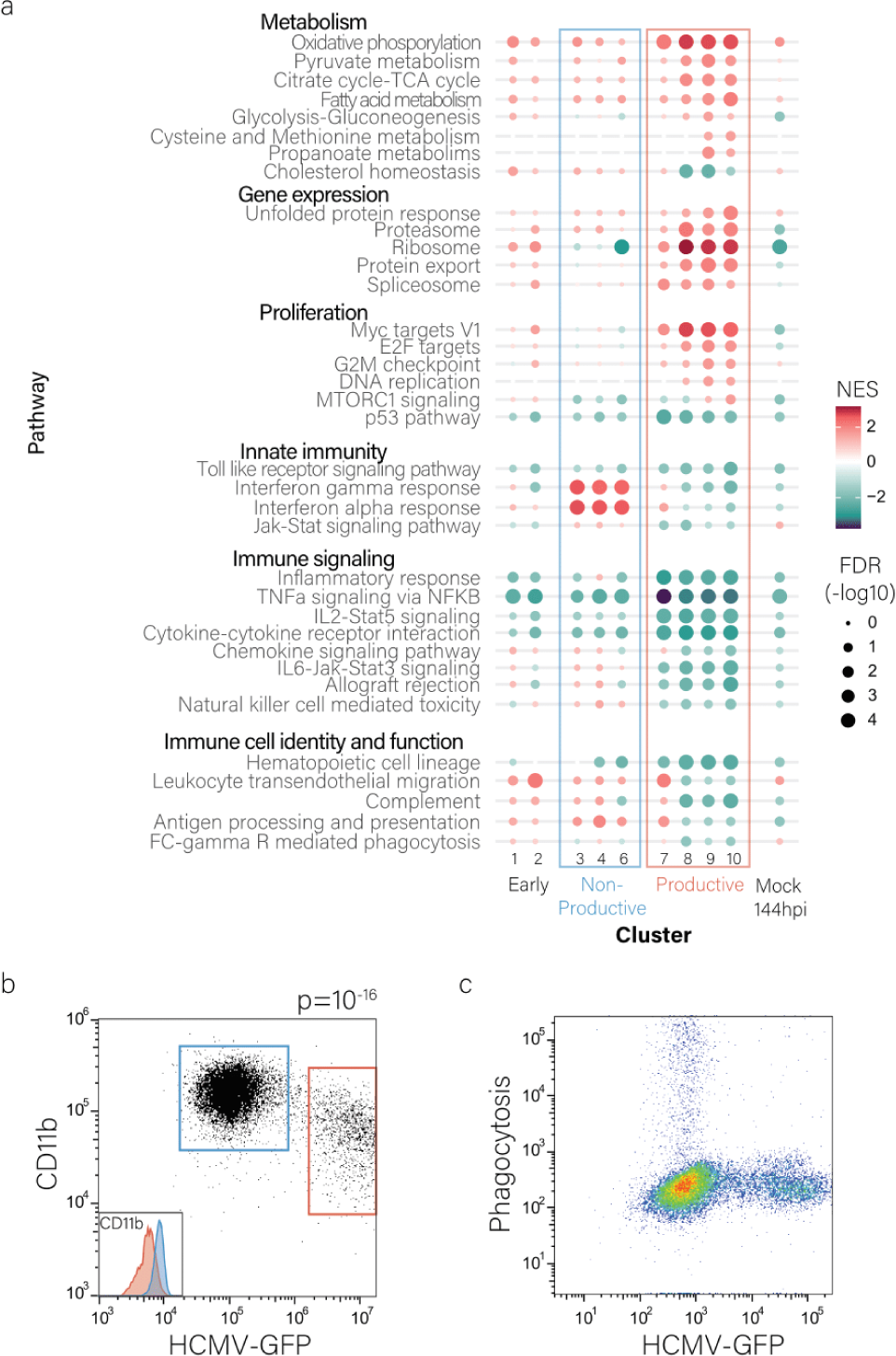
Effects of HCMV infection on cellular pathways and macrophage identity and function. a. Summary of gene set enrichment analysis (GSEA) of differentially expressed genes between each of the scRNA-seq clusters and mock-infected cells from 5 hpi using annotated KEGG and Hallmark pathways. Uninfected MDMs at 144 hpi are shown as control for changes occurring with time in culture regardless of infection. NES- normalized enrichment score. FDR- false discovery rate. b. Flow cytometry of CD11b surface levels in HCMV-GFP infected M-CSF-induced MDMs. The blue and red gates mark the non-productive and productive MDMs, respectively, as measured by GFP level. The inset in the bottom left shows the distribution of CD11b levels in the two populations. p value of the decrease in the mean fluorescence intensity in GFP-bright MDMs on the measured single cells is calculated using one-tailed t-test. Representative of two independent replicates is shown. c. Phagocytic activity measured by flow cytometry of HCMV- GFP infected M-CSF-induced MDMs. Replicate data and statistics are shown in Extended Data Fig. 6c.

Several pathways were suppressed in productive MDMs, including some which are associated with immune signaling and some which are associated with common functions of macrophages, such as antigen presentation and phagocytosis . Furthermore, genes related to hematopoietic cell lineage were also suppressed in productive MDMs (Fig. 6a). We further performed RNA-seq on infected BAL cells, enriched for macrophages, and sorted into GFP-bright and GFP-dim as well as on mock-infected cells. We show that indeed similar pathways related to immune signaling as well as macrophage identity and function are significantly suppressed in productively infected BAL macrophages (Extended Data Fig. 6a). In the non-productively infected cells the most noticeable changes, as in MDMs is strong induction of ISGs.

Since transcriptional signatures pointed to disruption of macrophage identity and function in productively infected MDMs, we next examined the surface levels of CD11b, a prominent macrophage marker, in MDMs induced by either PMA or M-CSF, and found reduced surface levels in productively infected cells (Fig. 6b and Extended Data Fig. 6b). Analysis of phagocytosis was done in M-CSF-induced MDMs, which regardless of infection exhibited more robust phagocytosis than PMA-induced MDMs, and revealed dramatically reduced phagocytosis activity in productively infected cells (Fig. 6c and Extended Data Fig. 6c). These results show, both on the transcriptional and the functional levels, that productive HCMV infection disrupts macrophage identity and function.

### Viral gene expression in HCMV infected MDMs

We next examined viral gene temporal expression along productive and non-productive infection. For the productive MDMs we binned the cells according to % viral reads to reflect progression along infection. In order to look at dynamic expression of viral genes, we calculated the relative expression pattern, out of total viral reads in a given bin. This was followed by hierarchical clustering, revealing a temporal expression cascade in productive MDMs that is similar to that seen in infection of fibroblasts ^28^ (Fig. 7a, left heatmap). It is important to note that this analysis reflects relative expression patterns, i.e. how each viral gene is expressed relative to all other viral genes. At the absolute level, almost all viral genes increase as infection progresses (Extended Data Fig. 7a). We next examined gene expression in the non-productive cells. We noticed that despite viral transcript levels being low throughout infection, intriguingly, they seem to increase with time (Fig. 7b), and this relative increase in viral gene expression was not observed in infected monocytes (Extended Data Fig. 7b). Although we computationally monitor and subtract reads that can originate from low level cross contamination between cells (by subtracting the background signal we capture from empty wells, see methods section), we wanted to exclude the possibility that this apparent increase in viral reads originates from low levels of cross contamination from the productive cells that were processed simultaneously with the non-productive MDMs and at late time points express extremely high levels of viral transcripts. To address this issue and to validate that the increase in viral reads is authentic, we conducted another experiment in which at 72 and 144 hpi only GFP-dim cells were processed separately allowing us to avoid productive cells and potential cross-contamination. This more stringent analysis further demonstrated a small increase in viral transcript levels along time post infection in non-productive MDMs (Extended Data Fig. 7c). We also examined if there are changes in viral gene expression in non-productive cells along infection. Besides UL123,which similarly to productive MDMs, was detected mainly at an early time point, we observed broad gene expression with no clear kinetic program (Fig. 7a, right heatmap). Overall, these results indicate that non-productive MDMs express diverse viral transcripts and compared to monocytes are more permissive for maintaining low levels of viral gene expression.

**Figure 7.**
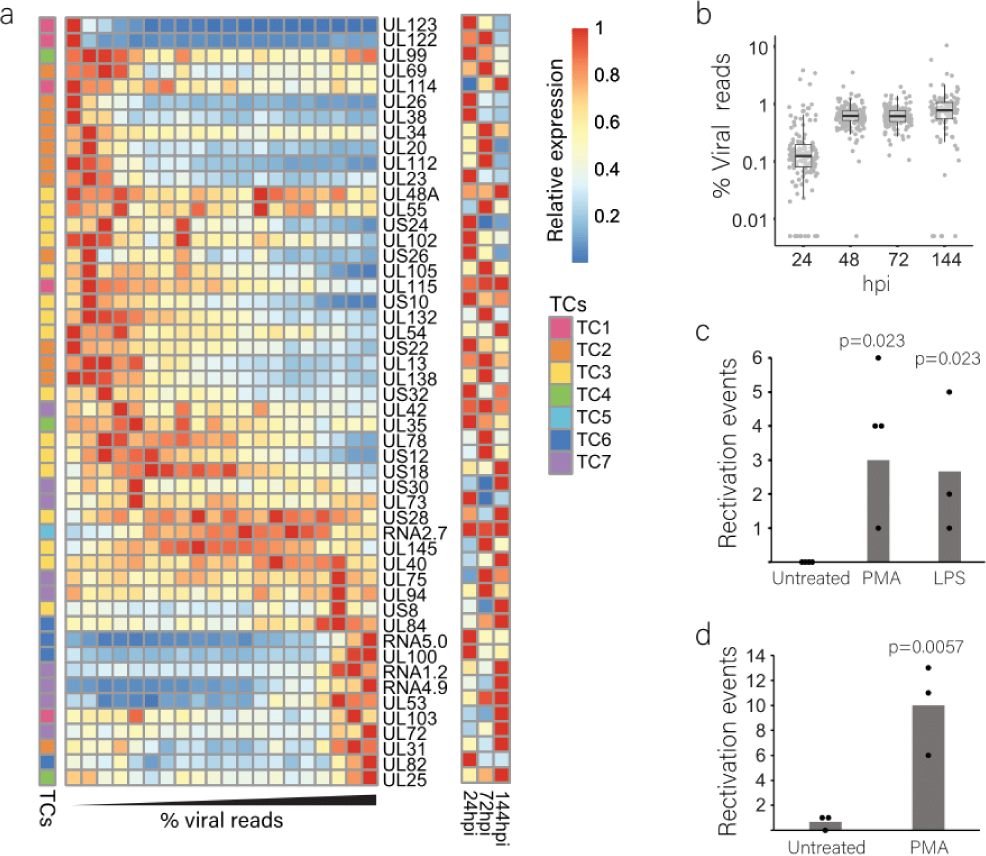
Non-productive MDMs exhibit mild elevation of gene expression with time and can reactivate to produce viral progeny. **a**. Heat map depicting normalized relative levels of the 50 top relatively expressed viral transcripts in productive MDMs (left heatmap) and in non-productive MDMs (right heatmap). Productive MDMs are binned and staged by percentage of viral reads. The expression levels in the heatmap are relative to the highest level of each transcript. Viral genes are ordered based on hierarchical clustering according to the productive MDMs. Annotations according to temporal classes ^28^ are shown to the left of the heatmaps. **b**. Percentage of viral reads in single non-productive MDMs along infection. **c**. Number of reactivation events in HCMV infected, GFP-dim sorted, M-CSF-induced MDMs following different treatments. Points show measurements of biological replicates. p-value of the increase in reactivation events upon stimuli, as calculated by one tailed t-test on the replicates, is indicated. **d**. Number of reactivation events in IFN pre-treated, HCMV infected, PMA-induced MDMs either left untreated or treated again with PMA. Points show measurements of biological replicates. p-value of the increase in reactivation events upon stimuli, as calculated by one tailed t-test on the replicates, is indicated.

### Non-productive MDMs can reactivate and release infectious virus

The observation that viral gene expression is maintained at higher levels in non-productive macrophages compared to monocytes, indicates a less repressed state in the former. Furthermore, in culture, these cells survived for relatively long periods (Extended Data Fig. 1c) and also *in-vivo* macrophages can be long lived cells ^16^. We therefore reasoned that non-productive MDMs could also potentially undergo reactivation. To explore this possibility, we infected both PMA and M-CSF induced MDMs, sorted the GFP-dim cells at 4 dpi, and at 7 dpi added either PMA or LPS as triggers for reactivation. We did not observe any reactivation without additional stimuli, attesting that our sorting indeed singled out only non-productive MDMs. However, both treatments resulted in reactivation events, forming GFP-positive plaques when co-cultured with fibroblasts (Fig. 7c, Extended Data Fig. 7d and 7e), illustrating that upon a stimuli, HCMV has the potential to reactivate from non-productive MDMs.

The complete elimination of productive cells following pre-treatment with IFN, brings up a question whether cells that are exposed to IFN prior to encountering the virus, are not infected, abortively infected or infected in similar manner to the non-productive MDMs we have characterized here, in which case they can later on reactivate. To test whether the last scenario is occurring, we infected IFN pre-treated MDMs with HCMV, and tested their ability to reactivate. As mentioned above, no productively infected cells could be detected following IFN pre-treatment, and at 7 dpi PMA was added as a trigger of reactivation. While there was mostly no reactivation without additional stimuli, indicating that infection in MDMs pre-treated with IFNs was indeed mostly non-productive, few reactivation events did occur, in line with previous findings ^43^, and PMA led to more robust reactivation of the virus accompanied by release of infectious progeny (Fig. 7d and Extended Data Fig. 7f).

These results show that at least in these culture models, MDMs can be infected in a latent-like infection, in which viral gene expression is repressed, but the viral genome retains the ability to reactivate upon an additional stimulation. HCMV infection following exposure to IFNs may further lead to a similar latent-like infection that can also result in reactivation.

## Discussion

When HCMV infects a cell, two outcomes are possible; a productive infection including propagation of the virus and release of infectious virus, or non-productive infection in which the viral genome is present in the cell but fails to propagate or produce infectious virus. Non-productive infection can potentially mean latent infection, if the viral genome is maintained and retains the ability to reactivate and complete productive infection. In order to delineate, in an unbiased manner, the determinants contributing to HCMV infection outcome, we comprehensively analyzed, by scRNA-seq, the progression of HCMV infection in monocytes and macrophages, which are considered to support latent and lytic infection, respectively.

In monocytes we found very low levels of viral transcripts in all cells, throughout the infection time course assayed, as was previously reported ^26–28^. Interestingly, infection of monocyte-derived macrophages (MDMs) resulted in both productive and non-productive infections. This was reflected in two distinct populations that differed in the levels of the viral-derived GFP expression, viral DNA replication and production of infectious progeny. Notably, MDMs exhibiting low virus-derived GFP levels harbored viral DNA indicating that they are not bystander cells and that they were infected. By applying scRNA-seq in a dense time course along infection we characterize these two distinct populations, showing low and high viral transcript loads. Productive cells exhibited a clear temporal viral gene expression cascade which resembles the kinetic viral gene expression in infected fibroblasts. In non-productive cells we detected low-level, broad viral gene expression, as has been reported previously for latent infection of monocytes ^26–28^. Importantly, no clear distinct populations could be detected at early times of infection, indicating there were no major differences prior to infection within the MDM population that can readily explain the different outcomes.

We found that the unequivocal factor associated with the earliest split of productive cells is viral gene expression and specifically the expression of immediate early genes and the onset of early gene expression. Our analysis revealed that the two immediate early genes, UL123 (IE1) and UL122 (IE2), are distinctly expressed at the split of productive and non-productive cells. Previously, ectopic expression of these genes in THP1 cells was shown to be insufficient to induce productive infection ^44^, which was explained by additional barriers. We show that overexpression of UL123 or UL122 but not a late viral gene, in THP1-derived macrophages results in much more cells being productively infected. The notable difference between the reports are the use of THP1 monocytic cells versus the use of differentiated THP1 cells. This suggests that in THP1-derived macrophages, these additional barriers are lower, allowing to reveal that sufficient expression of the IE genes is a major threshold for the establishment of productive infection. In addition, our findings imply that what defines the critical threshold of IE genes is their capacity to induce early genes. This possibly points to the ability to commence substantial viral DNA replication as the genuine step that determines infection outcome. The lack of a clear cellular gene expression signature that is associated with the initial increase in viral gene expression levels, indicates that in this experimental set-up, infection outcome is not dictated by variations in the cellular transcriptome and is likely mostly determined by the amount of infectious virus entering the cell ^29^, which will have significant consequences on the initial induction of viral gene transcription or possibly by variability between virion particles ^30^.

The defining signature of non-productive MDMs is the expression of ISGs. A similar feature was also shown in HSV-1 infected cells ^45^. Examination of viral transcript levels and expression levels of ISGs on a single cell level, generally showed that cells either elevate viral genes or elevate ISGs, thus seemingly defining two mutually exclusive scenarios. However, detailed examination of the trajectories along infection in both MDMs and fibroblasts strongly points out that productive infection can be established in cells that initially induced ISGs, which means that they may go through a stage of induction of ISGs and low viral transcript levels, before becoming productively infected. This illustrates that the induction of ISGs following infection does not deterministically define infection outcome, although it may account for delayed induction of viral gene expression. Indeed, addition of interferons to MDMs or fibroblasts immediately following infection did not affect the proportion of cells succumbing to productive infection.

On the other hand, we demonstrate that intrinsic expression levels of ISGs, independently of infection, influence infection outcome. Intrinsic levels of ISGs negatively correlate with permissiveness of cells to productive infection when comparing monocytes, MDMs and fibroblasts. Furthermore, differentiation of monocytes to macrophages as well along the hematopoietic lineage which are known to be associated with permissiveness to HCMV productive infection, is also associated with a decrease in intrinsic levels of ISGs. A connection between intrinsic expression of ISGs and differentiation was noted previously by Wu et al. ^46^ and intrinsic expression of select ISGs was shown to be functionally relevant in resistance of embryonic stem cells to infection by several RNA viruses ^46^. Our results suggest that this link between intrinsic expression of ISGs and differentiation, could be a major contributing factor to the cell-type dependent infection outcome of HCMV and potentially other herpesviruses.

Beyond explaining cell type differences, this effect of intrinsic levels of ISGs could be relevant to the establishment of latency. Notably, we show that interferon treatment prior to infection of both fibroblasts and MDMs almost completely abrogated productive infection. We further show that this treatment completely blocks viral DNA replication and acts at early stages of infection. These results support the notion that the levels of certain ISGs, at the time of infection, act to repress viral gene expression thus preventing productive infection. Similar findings were previously provided for murine CMV infection of fibroblasts and endothelial, where it was further shown that IFN-mediated silencing of the CMV genome is reversible ^43^. This effect of IFNs may have implications for infection in an *in-vivo* setting in which a substantial fraction of cells are not infected. In that case, the induction of IFNs by infected cells would affect neighboring uninfected cells by inducing ISG expression and subsequently shifting their fate towards non-productive infection, potentially aiding the virus to seed latent reservoirs, and perhaps also prolonging infection in case of reversion of silencing. Indeed we show that IFN pre-treated MDMs, which are infected non-productively, are able, upon stimulation, to reactivate, demonstrating that IFN exposure may lead to a latent-like infection in MDMs. Future work will be needed to delineate which of the ISGs contribute to blocking productive infection, and whether they are a central determinant in dictating cell-type dependent differences in permissiveness to productive infection and potentially establishment of latency.

Reactivation of HCMV from latency is a substantial clinical burden in immunocompromised individuals and may cause serious and often life-threatening disease ^47–49^. The only characterized HCMV latent reservoir to date is the hematopoietic cells in the bone marrow ^50–52^, however, the risk of HCMV reactivation following transplantations, depending on whether the donor and/or the recipient are carriers of HCMV, suggests that latent reservoirs exist also in tissues ^11^. For many years tissue resident macrophages were considered to be short lived and be replenished regularly by circulating monocytes. However, mounting evidence in recent years indicates that self-maintaining resident macrophage populations reside in all mammalian organs and can sustain for years ^16, 53, 54^. Our analysis demonstrates that macrophages can be non-productively infected, they can be viable for weeks in culture, carry viral genomes and harbor low levels of viral transcripts. Importantly, we show that in culture these viral genomes can reactivate, suggesting that infected macrophages can potentially be latently infected. Interestingly, HCMV reactivation was demonstrated in alveolar macrophages from healthy HCMV carriers ^55^. Another support for the idea that macrophages may serve as a latent HCMV tissue reservoir comes from an observation made 25 year ago that macrophages carry latent MCMV following clearance of an acute infection ^56^.

Macrophages are found in most tissues and are the first responders to microbial invasion, thus are an important component of the immune system. Our analysis of productively infected macrophages demonstrates a dramatic effect on macrophage identity and function. This analysis complements previous work that examined the effects of HCMV infection on monocytes ^8, 57–60^ and aligns with a recent study by Baasch et al. that showed productive MCMV infection of murine bone marrow-derived macrophages as well as alveolar macrophages *in vivo* reprograms macrophage identity and disrupts their function, leading to higher susceptibility to secondary lung infection in mice ^61^. We show here that productive HCMV infection leads to similar consequences in human monocyte-derived macrophages, disrupting macrophage identity and dramatically impairing phagocytosis. Thus, primary HCMV infection of tissue macrophages could interfere with their immune functions, and this may lead to increased susceptibility to additional infections.

All together, our results demonstrate that HCMV infection of macrophages can result in different outcomes, either productive infection or non-productive infection, a split which we show is mainly dictated by the ability of the virus to express its two major immediate early genes. We further reveal that the intrinsic level of ISGs greatly influences infection outcome and may explain the tendency of HCMV to establish productive infection in differentiated cells.

Finally, we show that productive infection perturbs macrophage function while non-productive infection is associated with repressed viral gene expression, and maintenance of viral genomes which has the potential to reactivate. These findings highlight macrophages as prospective critical players during HCMV infection both in pathogenesis at the time of acute infection, as well as potential novel latent reservoirs in tissues.

## Methods

### Cell culture and virus

Primary CD14+ monocytes were isolated from fresh venous blood, obtained from healthy donors, using Lymphoprep (Stemcell Technologies) density gradient followed by magnetic cell sorting with CD14+ magnetic beads (Miltenyi Biotec). Monocytes were cultured in X-Vivo15 media (Lonza) supplemented with 2.25 mM L-glutamine at 37°C in 5% CO2, at a concentration of 1-2 million cells/ml in non-stick tubes to avoid differentiation.

Where indicated, primary monocytes were treated with the following treatments immediately following isolation and plating: PMA 50ng/ml for 3 days, GM-CSF and IL-4 1000U/ml each for 6 days with cytokine replenishing after 3 days, M-CSF 20ng/ml for 6days with cytokine replenishing after 3 days, TNFɑ 10ng/ml for 6 days. All treatments were done in Roswell Park Memorial Institute (RPMI) with 10% heat-inactivated fetal bovine serum (FBS), 2 mM L-glutamine, and 100 units/ml penicillin and streptomycin (Beit-Haemek, Israel) and MDMs were grown in this media following treatment. Primary human foreskin fibroblasts (ATCC CRL-1634) were maintained in DMEM with 10% FBS, 2 mM L-glutamine, and 100 units/ml penicillin and streptomycin (Beit-Haemek, Israel). Interferon treatment was done with a combination of 550 U/ml IFNα, 700 U/ml IFNβ and 500 U/ml IFNγ. Ruxolitinib treatment was done at a concentration of 4μM. THP1 cells, purchased from ATCC (TIB-202), were grown in RPMI media with 15% heat-inactivated fetal bovine serum (FBS), 2 mM L-glutamine, and 100 units/ml penicillin and streptomycin (Beit-Haemek, Israel). Kasumi-3, purchased from ATCC (CRL-2725) were grown in RPMI media with 20% heat-inactivated fetal bovine serum (FBS), 2 mM L-glutamine, and 100 units/ml penicillin and streptomycin (Beit-Haemek, Israel). Differentiation of THP1 and Kasumi-3 cells was done by adding PMA 50ng/ml for 2 days. Infection and growth following infection were done in the media containing 10% FBS.

BronchoAlveolar Lavage fluid was obtained from routine bronchoscopies. 15-40ml were obtained from each sample and diluted by adding an equal volume of RPMI media with 10% heat-inactivated fetal bovine serum (FBS), 2 mM L-glutamine, and 100 units/ml penicillin and streptomycin (Beit-Haemek, Israel). Diluted BAL fluid was transferred through a 70uM filter and plated on a tissue culture dish. Adherent cells were washed thoroughly prior to infection.

The TB40E strain of HCMV, containing an SV40-GFP tag, was described previously ^17^. Virus was propagated by adenofection of infectious bacterial artificial chromosome (BAC) DNA into fibroblasts as was previously described ^62^. Viral stocks were concentrated by centrifugation at 26000xg, 4°C for 120 minutes. Infectious virus yields were assayed on primary fibroblasts.

### Infection and reactivation procedures

Infection was performed by incubating cells with the virus for 3 hr, washing twice and supplementing with fresh media. Initial experiments with primary non-differentiated and differentiated monocytes as well as the single cell experiments of monocytes and MDMs were done at MOI=10 to ensure infection of all the cells. Since this MOI is based on quantification of infectious particles in fibroblasts it is likely effectively lower in monocytic cells. The single cell experiments in BAL macrophages were done in MOI=5, and all other experiments were done in MOI=2. For reactivation assays, GFP-dim infected MDMs were sorted at 4dpi and plated in 96- well plates (10,000 cells per well), IFN-pretreated MDMs were used following infection without further sorting. At 7dpi cells were either treated, as indicated, with 50ng/ml PMA (Sigma), or 500ng/ml LPS (Sigma) or left untreated for 3 days and then the media was replaced with fresh untreated media. At 14dpi, 7 days post treatment, GFP positive cells were counted on a fluorescent microscope. At 18dpi 7500 fibroblasts per well were seeded with the MDMs and plaque formation was followed and imaged for a period of 14 days.

### Flow cytometry and sorting

Cells were analyzed on a BD Accuri C6 or a BD LSRII and sorted on a BD FACSAria III. All analyses and figures were done with FlowJo. All histograms were plotted with modal normalization.

### Measurement of viral progeny

Fibroblasts were infected with known volumes of supernatants taken from infected MDM or monocyte cultures. Number of GFP positive fibroblasts were measured at 2dpi by flow cytometry and the infectious particles per volume was calculated.

### Measurement of viral genomes by digital PCR

DNA was extracted from cell pellets in a 1:1 mixture of lysis solutions A (100 mM KCl, 10 mM Tris–HCl pH 8.3, and 2.5 mM MgCl2) and B (10 mM Tris–HCl pH 8.3, 2.5 mM MgCl2, 0.25% Tween 20, 0.25% Non-idet P-40, and 0.4 mg/ml Proteinase K), for 60 min at 60°C followed by a 10 min 95°C incubation, according to the description in Roback et al. ^63^. Measurement of viral DNA was done using the QX200 droplet digital PCR system (Bio-Rad), using FAM labeled HCMV primer and probe: Human CMV HHV5 kit for qPCR using a glycoprotein B target (PrimerDesign); and HEX labeled RPP30 copy number assay for ddPCR (Bio-Rad), as previously described ^64^. Analysis was done using the Bio-Rad QX software.

### Microscopy

Imaging was performed on an Olympus Ix73 inverted fluorescent microscope using a X4 or a X10 objective and an Olympus DP73 camera or on an AxioObserver Z1 wide-field microscope using a X20 objective and Axiocam 506 mono camera.

### Single cell RNA sequencing library preparation

For primary monocytes and MDMs, single-cell sorting and library preparation were conducted according to the massively parallel single-cell RNA-seq (MARS-seq) protocol, as previously described ^65^. In brief, single cells were FACS sorted into wells of 384-well capture plates containing 2μl of lysis buffer and reverse transcription (RT)-indexed poly(T) primers, thus generating libraries representing the 3’ end of mRNA transcripts. Eight empty wells were kept in each 384-well plate as a no-cell control during data analysis. Immediately after sorting, each plate was spun down to ensure cell immersion into the lysis solution, snap-frozen on dry ice, and stored at -80°C until processed. Barcoded single-cell capture plates were prepared with a Bravo automated liquid handling platform (Agilent). For generation of the RNA library, mRNA from single cells was converted into cDNA in the capture plates and pooled by centrifugation to VBLOCK200 reservoir (Clickbio). The pooled sample was linearly amplified by T7 in vitro transcription, and the resulting RNA was fragmented and converted into a sequencing-ready library by tagging the samples with pool barcodes and Illumina sequences during ligation, RT, and PCR. Each pool of cells was tested for library quality, and concentration was assessed as described earlier ^65^.

For BAL cells, pools of 10,000 cells were prepared for scRNA-seq using Chromium Single Cell 3′ Gene Expression v3.1 kit according to the manufacturer’s instructions (10x Genomics).

### sc RNA-seq data analysis

Analysis of MARS-seq data was done with the tools described in Jaitin et al. and Paul et al. ^66, 67^. The reference was created from the hg19 and TB40E (NCBI EF999921.1) strain of HCMV. The transcription units of the virus were based on NCBI annotations, with some changes based on the alignment results. This includes merging several transcripts (taking into account that the library maps only the 3’ ends of transcripts) and adding some antisense transcripts. Read assignment to wells was based on the batch barcode (4 bp) and the well barcode (7 bp) and reads with low-quality barcodes were removed. The trimmed read (37 bp) was aligned with the reference using Bowtie 2 ^68^, and the counting of the reads is done based on unique molecular identifiers (UMIs, 8 bp). For each batch, the leakage noise level was estimated by comparing the number of UMIs in the eight empty wells to the total number of UMIs in the batch. The noise levels in all batches was lower than 1.5% . Wells with less than 500 or more than 5,000 expressed genes, or more than 20,000 reads were discarded. The number of wells that were used for further analysis is 3,226 .

For 10xGenomics data, we used Cell Ranger software (V6.1.2, 10x Genomics) with the default settings to process the FASTQ files with the references used in the analysis of the MARS-seq data. Cells with less than 500 or more than 10,000 expressed genes, or more than 30,000 reads were discarded. The number of wells that were used for further analysis is 21,839 .

All single cell analyses were done using Seurat V4.0.3 ^69^. To visualize single-cell datasets, we performed dimension reduction by Uniform Manifold Approximation and Projection (UMAP), based on both cellular and viral genes in figures 2 and 3 or only cellular genes in Fig. 4 and Extended Data Fig. 4). Clusters of MDMs were numbered based on % viral reads in the cluster, from lowest to highest.

### Calculation of viral and cellular gene expression levels in single cells

Expression of a single cellular gene or a group of cellular genes in single cells is calculated as the percentage of reads on this gene or group of genes out of the total cellular reads in a cell. % viral reads is calculated as the percentage of reads on all viral genes out of all reads in a cell. These numbers were used for coloring UMAP plots in figures 2d, 3c, 3f, 5a, Extended Data Fig. 3b and for plots in figures 3e, 5b, 5c, 5g, 5h, 5i, 5j, 7b, Extended Data figures 5b, 5f, 7b, 7c. The list of ISGs is the combined list of the following gene sets: Hallmark interferon alpha response, Hallmark interferon gamma response and Reactome interferon signaling. The list of Macrophage differentiation genes is based on the list of genes in the GO term regulation of macrophage differentiation (GO:0045649).

Relative gene expression level of a single viral gene or a group of viral genes is calculated as the percentage of reads on this gene or group of genes out of the total viral reads in a cell. These numbers were used for coloring the UMAP plot in Fig. 4c and for plots in Fig. 4b, and Extended Data Fig. 4b.

Subtraction of cross contaminating viral reads was done in the following way: for each plate the level of noise was estimated from the ratio between the average number of UMIs in 8 empty wells to the average number of UMIs in the plate. This number was used to calculate the fraction of viral UMIs that can be attributed to leakage between cells, and was subtracted from the number of viral reads in all cells of the plate.

### Bulk RNA-seq library preparation and alignment

For RNA-seq of THP1 cells, cells were collected with Tri-Reagent (Sigma-Aldrich), total RNA was extracted by phase separation and poly(A) selection was performed using Dynabeads mRNA DIRECT Purification Kit (Invitrogen) according to the manufacturer’s protocol. RNA-seq libraries were generated as previously described ^70^. mRNA samples of 4 ng were subjected to DNaseI treatment and 3’ dephosphorylation using FastAP Thermosensitive Alkaline Phosphatase (Thermo Scientific) and T4 PNK (NEB) followed by 3’ adaptor ligation using T4 ligase (NEB). The ligated products were used for reverse transcription with SSIII (Invitrogen) for first-strand cDNA synthesis. The cDNA products were 3’ ligated with a second adaptor using T4 ligase and amplified with 8 cycles of PCR for final library products of 200–300 base pairs. Raw sequences were first trimmed at their 30 end, removing the illumina adapter and polyA tail. Alignment was performed using Bowtie 1 ^71^ (allowing up to 2 mismatches) and reads were aligned to the human (hg19). Reads aligned to ribosomal RNA were removed. Reads that were not aligned to the genome were then aligned to the transcriptome.

For RNA-seq of BAL macrophages, pellets from cell sorting were lysed in RLT buffer (Qiagen) containing 40mM DTT. Due to the low number of cells, RNA-seq libraries were performed using a modified mcSCRB-seq ^72^ adapted for bulk RNA-seq. Briefly, RNA was isolated using SPRI beads, reverse transcription and template switching was performed with indexed poly(T) primers in the presence of polyethylene glycol, followed by amplification of the final library.

66-bp raw reads were aligned using STAR (V2.7.1) ^73^ to concatenation of the human (hg19) and the viral genomes (NCBI EF999921.1). Counting of reads per gene was done based on unique molecular identifiers (UMIs, 10 bp).

### Plasmids and lentiviral transduction

UL123, UL122, UL99 or mCherry-flag were cloned into pLVX-Puro-TetONE-SARS-CoV-2-nsp1- 2XStrep (kind gift from Nevan Krogan, UCSF) in place of the SARS-CoV-2-nsp1-2XStrep cassette. UL123 was also cloned into pLVX-EF1alpha-SARS-CoV-2-nsp1-2XStrep-IRES-Puro (Addgene plasmid #141367) in place of the SARS-CoV-2-nsp1-2XStrep cassette. The UL123, UL122, and UL99 coding sequence were amplified from cDNA of HCMV infected fibroblasts with the primers detailed below, containing flanking regions homologous to the vector. The mCherry-flag was amplified from an expression plasmid using the primers detailed below, containing flanking regions homologous to the vector. The plasmids were amplified with the primers detailed below. The amplified PCR fragments were cleaned using a gel extraction kit (Promega) according to the manufacturer’s protocol. mCherry-flag was cloned into the vector by restriction-free cloning ^74^ and the viral genes were cloned into the vectors using a Gibson assembly protocol ^75^.

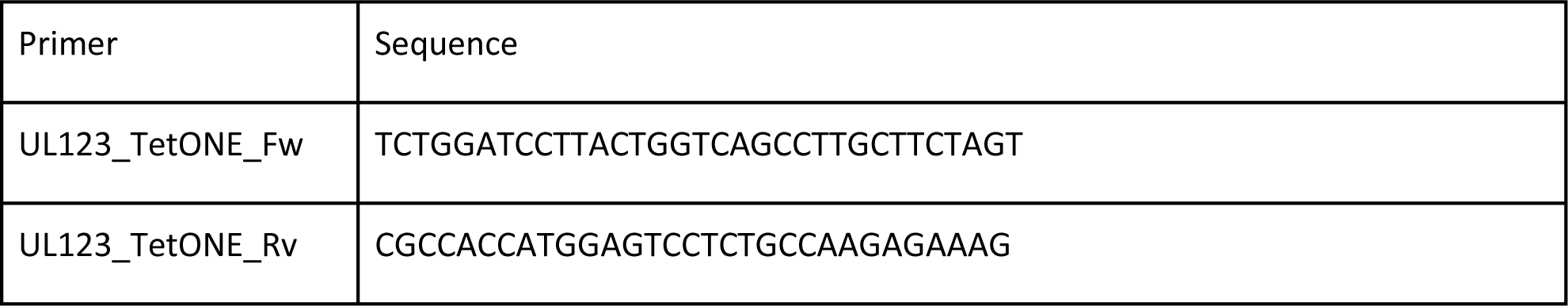
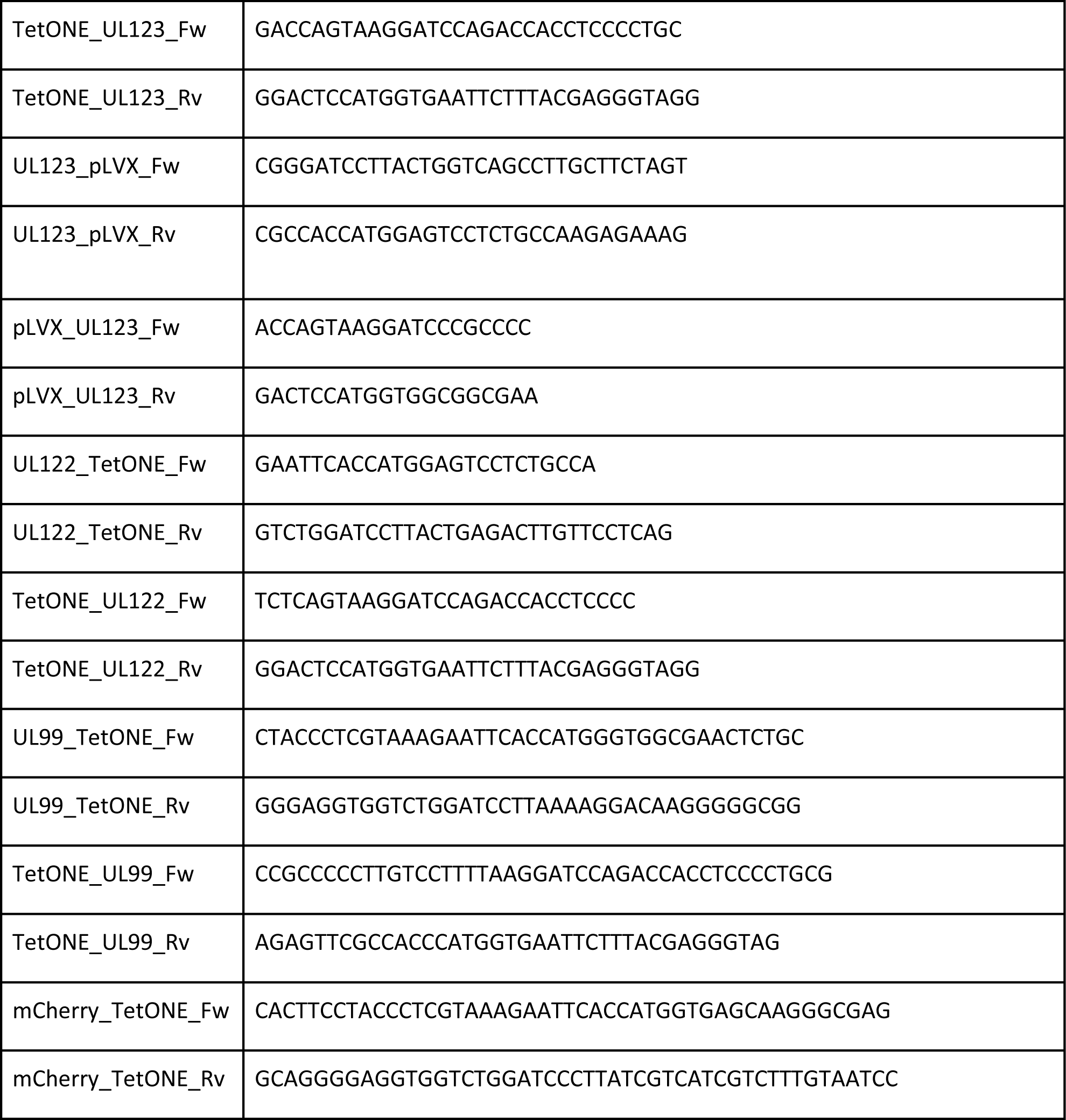

CRISPR KO was done using a CRISPR–cas9 system, with lentiCRISPR v2 plasmid (Addgene no. 52961). The following sgRNAs were cloned downstream of the U6 promoter: IRF9_1: 5′- AATTTAAGGAGGTTCCTGAG-3′; IRF9_2: 5′-ACAATTCCACAGGCCAGCCA-3′; STAT2_1: 5′- ATCATCTCAGCCAACTGGGT-3′; STAT2_2: 5′-AGCAACATGAGATTGAATCC-3′; IGSF8 (control): 5′- GCGGCAGCAGCGTGGGCCTGA-3’.

Lentiviral particles were generated by co-transfection of the expression constructs and 2nd generation packaging plasmids (psPAX2, Addgene#12260 & pMD2.G, Addgene#12259), using jetPEI DNA transfection reagent (Polyplus transfection) into 293T cells, according to the manufacturer’s instructions. 60 hours post transfection, supernatants were collected and filtered through a 0.45μm PVDF filter (Millex). THP1 cells were transduced with lentiviral particles by spinfection (800xg, 1hr) and were then puromycin-selected (1ug/mL) for 4 days. Puromycin was removed prior to subsequent differentiation. Induction of the inducible transgene was done by addition of 100ng/ml Doxycycline (Sigma) 24 hours prior to HCMV infection.Kasumi-3 cells were transduced with lentiviral particles by spinfection (800xg, 1hr). Since Doxycycline induction was not successful in our hands in Kasumi-3 cells, they have been transduced with a non-inducible version of UL123 expression plasmid and a control mCherry expression plasmid. To avoid long term effects of the transgene expression no selection step was done and the cells were treated with PMA two days after transduction followed by HCMV infection two days later.

### Quantitative real-time PCR analysis

For analysis of RNA expression, total RNA was extracted using Tri-Reagent (Sigma) according to the manufacturer’s protocol. cDNA was prepared using qScript cDNA Synthesis Kit (Quanta Biosciences) according to the manufacturer’s protocol. For analysis of DNA levels, cells were lysed in a 1:1 mixture of PCR solutions A (100 mM KCl, 10 mM Tris–HCl pH 8.3, and 2.5 mM MgCl2) and B (10 mM Tris–HCl pH 8.3, 2.5 mM MgCl2, 1% Tween 20, 1% Non-idet P-40, and 0.4 mg/ml Proteinase K), for 60 min at 60°C followed by a 10 min 95°C incubation. Real time PCR was performed using the SYBR Green PCR master-mix (ABI) on the QuantStudio 12K Flex (ABI).

The following primers (forward, reverse), spanning an exon-exon junction, were used for viral transcript quantification, for ISG transcript quantification, and the for the host gene Anxa5 that was used for normalization:

UL123: GCGCCAGTGAATTTCTCTTC, GTCCTGGCAGAACTCGTCA

UL22a: TTACTAGCCGTGACCTTGACG, CAGAAATCGAAGCGCAGCG

ISG15: TTTGCCAGTACAGGAGCTTG, TTCAGCTCTGACACCGACAT

MX1: ACCATTCCAAGGAGGTGCAG, TGCGATGTCCACTTCGGAAA

IFI6: TACACTGCAGCCTCCAACTC, AGTTCTGGATTCTGGGCATC

Anxa5: AGTCTGGTCCTGCTTCACCT, CAAGCCTTTCATAGCCTTCC

DNA was extracted from cell pellets or directly from adherent cells in a 1:1 mixture of lysis solutions A (100 mM KCl, 10 mM Tris–HCl pH 8.3, and 2.5 mM MgCl2) and B (10 mM Tris–HCl pH 8.3, 2.5 mM MgCl2, 0.25% Tween 20, 0.25% Non-idet P-40, and 0.4 mg/ml Proteinase K), for 60 min at 60°C followed by a 10 min 95°C incubation, according to the description in Roback et al. ^63^. The following viral primers (forward, reverse) were used to quantify viral DNA and a host gene (B2M) was used for normalization:

UL44: AGCAAGGACCTGACCAAGTT, GCCGAGCTGAACTCCATATT

B2M: TGCTGTCTCCATGTTTGATGTATCT, TCTCTGCTCCCCACCTCTAAGTWestern blot analysis

### Western blot analysis

Cells were lysed in RIPA buffer for 30 minutes on ice and then centrifuged at 14,000 xg for 20 min at 4°C. Samples were separated by 4 to 12% polyacrylamide Bis-Tris gel electrophoresis (Invitrogen), blotted onto nitrocellulose membranes, and immuno- blotted with the following primary antibodies: mouse anti-CMV IE1 and IE2 (abcam), mouse anti-PP28 (EastCoast Bio), rabbit anti-GAPDH (Cell Signaling Technology), rabbit anti- IRF9 (Cell signaling Technology), rabbit anti- STAT2 (Cell signaling Technology), mouse anti- actin (Sigma). The secondary antibodies were goat anti-rabbit–IRDye 800CW and goat anti-mouse–IRDye 680RD (LI-COR). Membranes were visualized in a LI-COR Odyssey imaging system.

### Correlation between cellular gene expression and viral gene expression levels

For calculating the Spearman correlation between cellular genes and viral expression levels, in each cell, the sum of viral reads was normalized to the total number of reads in the cell, and the number of reads of each cellular gene was normalized to the total number of cellular reads in the cell. The correlation was calculated across 313 and 310 cells on 5007 and 5606 genes for the 5 hpi and 12 hpi analyses, respectively. Genes with a total number of reads less than 45 (after normalization to 5,000 reads per cell) were not included.

### Differential expression and enrichment analysis

Differential expression analysis on bulk RNA-seq data was performed with DESeq2 (V1.22.2) ^76^ using default parameters, with the number of reads in each of the samples as an input. Differential expression analysis of the scRNA-seq was done using Seurat V4.0.3 ^69^

The log2 fold change values were used for enrichment analysis using GSEA (version 4.1, ^77^). For scRNA-seq data, only genes that were expressed in at least 20% of cells in both groups of cells in a comparison were used. For bulk RNA-seq, only genes that had at least 10 reads in each sample were used. The gene sets that were used were Hallmark and KEGG from MSigDB (version 7.4, 78).

### Viral gene expression heatmaps

Viral gene expression heatmaps depict the viral gene expression level either relative to total number of viral reads (figure 7a) or normalized to the total number of reads (figure S7a). The expression levels in the heatmap are relative to the highest level of each gene. Productive MDMs were ordered by their % viral reads. The cells were then grouped according to the number of viral reads per bin increasing from the lowest bin to the highest bin, to ensure a similar number of cells per bin. The heatmap shows the 50 viral genes with the top relative expression at their highest expressing bin. Non-productive cells were grouped according to the time post of infection. In each bin/time group, expression level of each viral gene was calculated as the total number of reads on the gene divided by the total number of viral reads.

### Marker staining and phagocytosis assay

Cells were stained in cold MACS buffer (PBS, 5% BSA, 2 mM EDTA). Cell staining was done using an APC-conjugated anti-human CD11b (#101211, Biolegend), PE-conjugated anti-human CD11c (#333149, BD), or FITC-conjugated anti-human CD14 (#130-080-701, Miltenyi). Phagocytosis was assayed using a phagocytosis kit (#601490, Cayman chemical) according to the manufacturer’s instructions.

### Graphics

Figure 2a was created with BioRender. Flow cytometry figures were created with FlowJo. Figure 4a was created with the R package ggalluvial. Figure 5d was created with R package sankey.

### Ethics statement

All fresh peripheral blood samples were obtained after approval of protocols by the Weizmann Institutional Review Board (IRB application 92–1) and following informed consent from the donors. The study using BronchoAlveolar Lavage (BAL) fluid samples was approved by a local research ethics committee in accordance with the Declaration of Helsinki (HMO-0704-20). Written informed consent to retain BAL fluid was obtained from patients undergoing bronchoscopy.

### Data availability

Sequencing data have been deposited in GEO under accession code GSE189581.

## Acknowledgements

We thank Nir Drayman, Roni Winkler, and the members of the Stern-Ginossar lab, for critical reading of the manuscript. We thank Dr. Keren Bahar Halpern and Dr. Shani Ben-Moshe for assistance with mcSCRB-seq and 10xGenomics library preparation. We thank Eain A Murphy for the TB40E-GFP virus strain. We thank the Weizmann flow cytometry unit for technical assistance. This study was supported by a European Research Council consolidator grant to N.S-G (CoG-2019-864012). N.S-G is a member of the European Molecular Biology Organization (EMBO) Young Investigator Program.

---

**Extended Data Figure 1.**
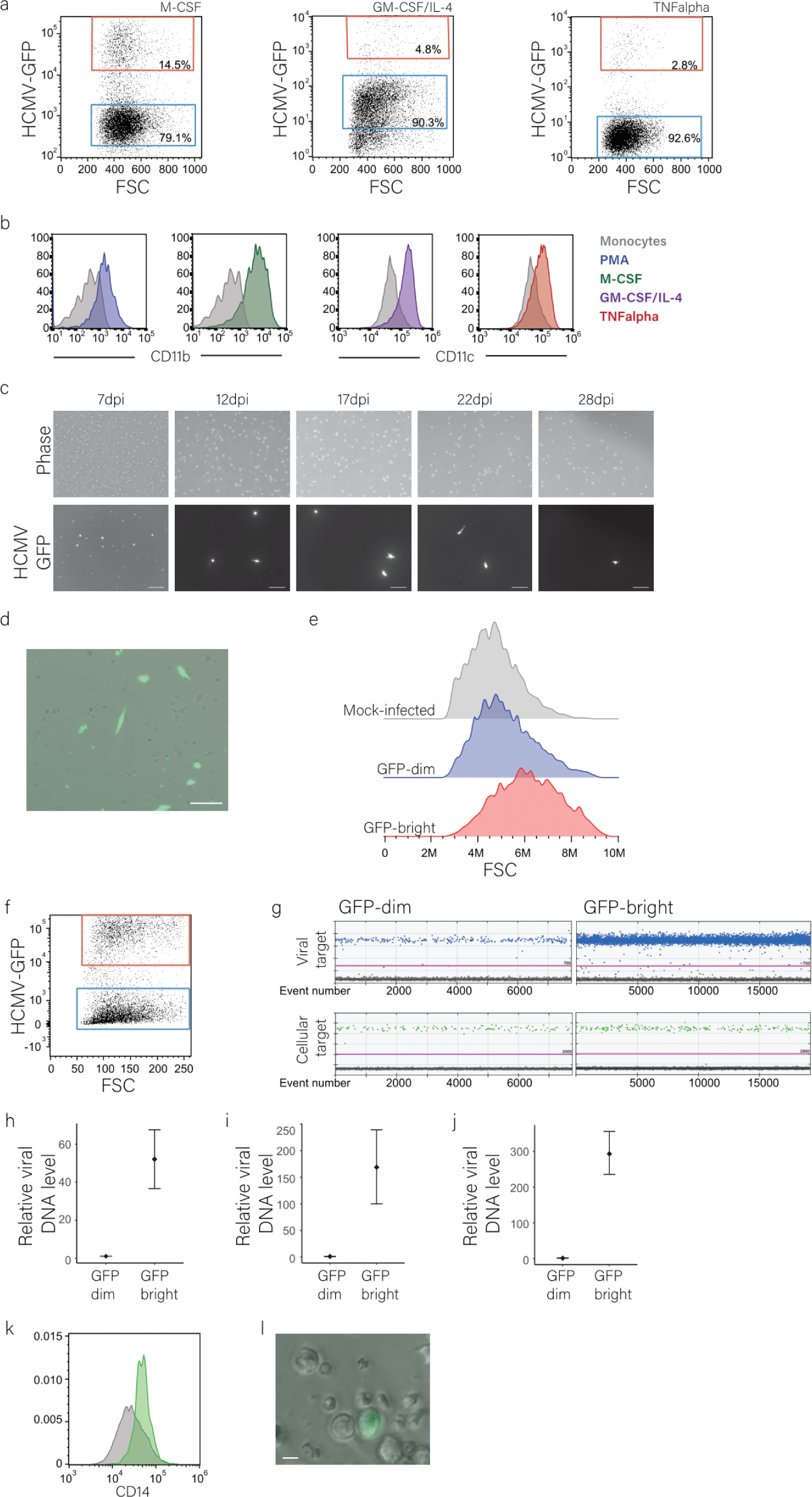
Productive and non-productive infection in monocyte derived macrophages. **a**. Flow cytometry analysis of primary CD14+ monocytes treated with the indicated treatments and infected with HCMV-GFP. Analysis was performed at 3dpi.Blue and red gates mark the GFP-dim and GFP-bright populations, respectively, and their percentage is noted. **b**. Flow cytometry analysis of CD11b surface expression in PMA and M-CSF treated monocytes and of CD11c surface expression in GM-CSF/IL-4 and TNF alpha treated monocytes compared to primary CD14+ monocytes. **c**. Phase-contrast and GFP microscopy images of PMA-induced MDMs infected with HCMV-GFP at different days post infection. Scale bars are 200μm. **d**. Microscopy image of PMA-induced MDMs at 3 dpi with HCMV-GFP. Scale bar is 100μm. **e**. Flow cytometry analysis of PMA-induced MDMs infected with HCMV-GFP at 3 dpi showing the size, measured by forward scatter (FSC), of mock-infected, GFP-dim and GFP-bright cells. **f**. Flow cytometry plot showing the gates used to sort GFP-bright (red gate) and GFP-dim (blue gate) HCMV-GFP infected PMA-induced MDMs at 4 dpi. **g**. digital droplet PCR (ddPCR) results of GFP-dim and GFP-bright HCMV-GFP infected MDMs at 4 dpi. Upper panel shows detection of viral DNA, lower panel reflects detection of cellular genomes. The magenta line marks the threshold. **h-j**. Measurements of viral genomes by ddPCR from GFP-dim and GFP-bright FACS-sorted populations of monocytes treated with M-CSF (**h**), GM-CSF/IL-4 (**i**) or TNFa (**j**). ddPCR graph is a representative of two biological replicates and shows the mean and 95% CV of poisson distribution. **k.** Flow cytometry analysis of CD14 surface expression in BAL cells enriched for macrophages. Stained and unstained cells are in green and gray, respectively. **l.** Microscopy image of HCMV-GFP infected BAL cells at 2dpi. Scale bar is 10μm.

**Extended Data Figure 2.**
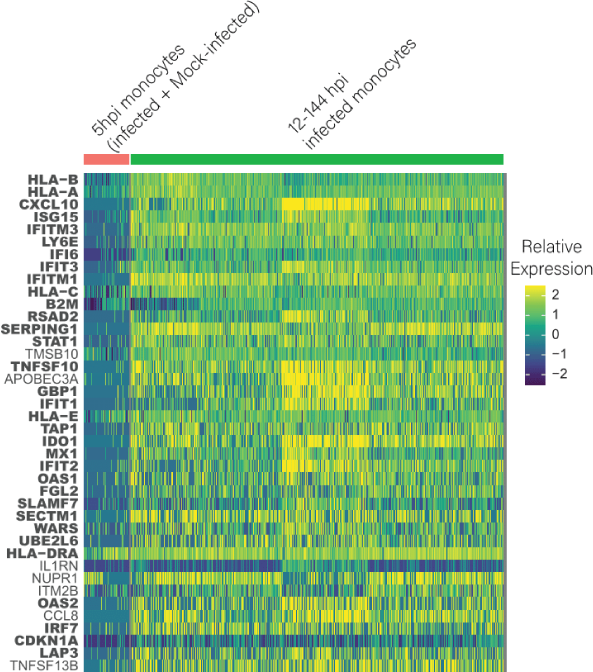
Infection in monocytes upregulates mainly interferon stimulated genes Heatmap depicting the 40 most significantly increased genes between 5hpi infected and mock-infected monocytes and 12hpi-144hpi infected monocytes. Interferon stimulated genes are highlighted in bold. Z-scores of the log of the expression values are shown.

**Extended Data Figure 3.**
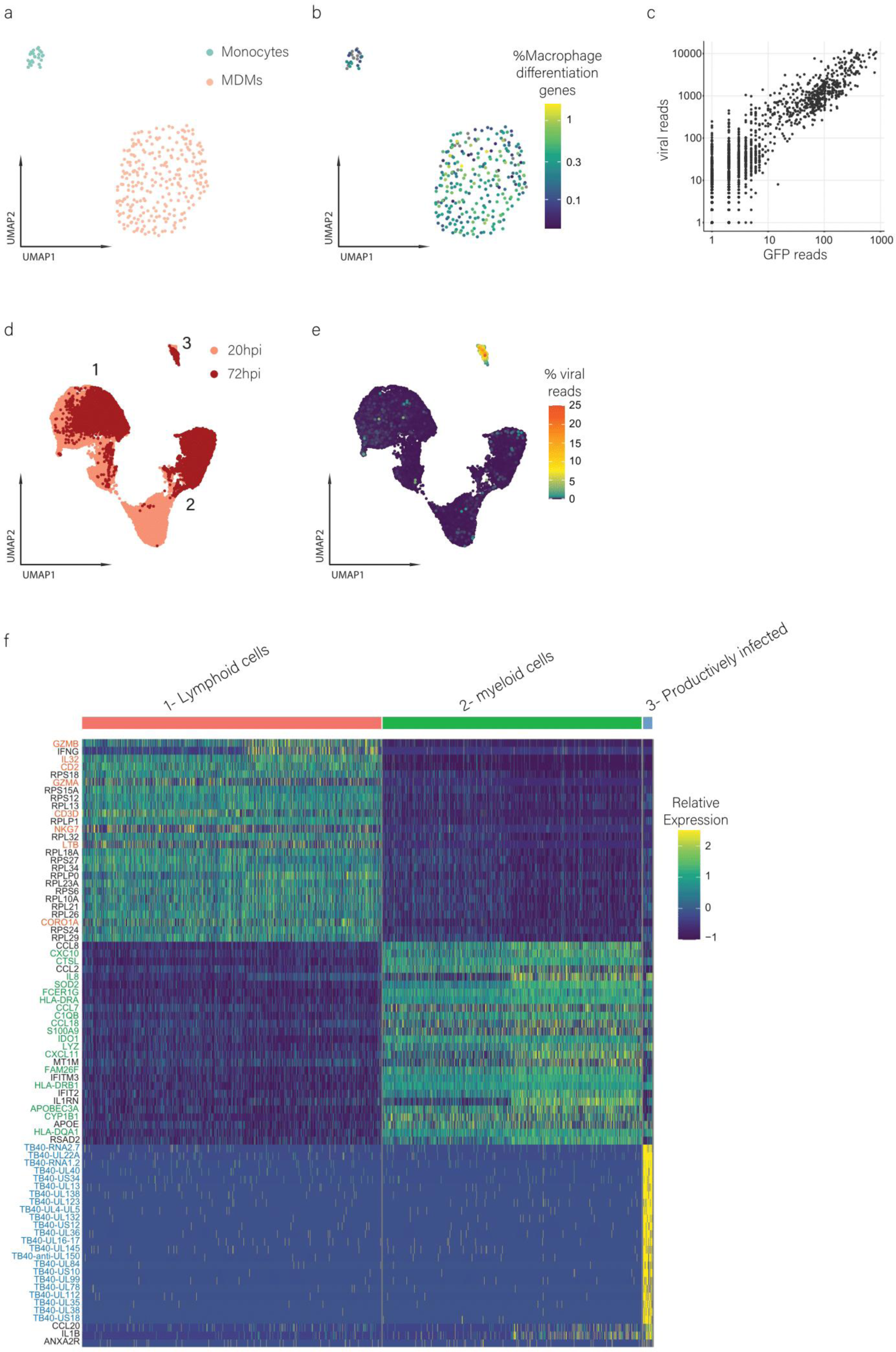
General features of single infected MDMs and BAL cells **a**. Projection of 283 mock-infected monocytes and MDMs at 5 hpi colored by cell type. **b**. Projections as in **a**, colored by the expression level of macrophage differentiation genes. **c**. Levels of viral reads versus GFP reads in single HCMV-GFP infected MDMs. **d.** Projection of 21,839 HCMV-infected BAL cells from 20 and 72 hpi colored by time post infection. Analysis was performed based on both cellular and viral transcripts. **e.** Projection as in d, colored by percentage of viral reads per cell. **f.** Heatmap depicting the most significantly variable genes between the different groups of BAL cells. Lymphoid, myeloid and viral genes are labeled in red, green and blue, respectively. Z-scores of the log of the expression values are shown.

**Extended Data Figure 4.**
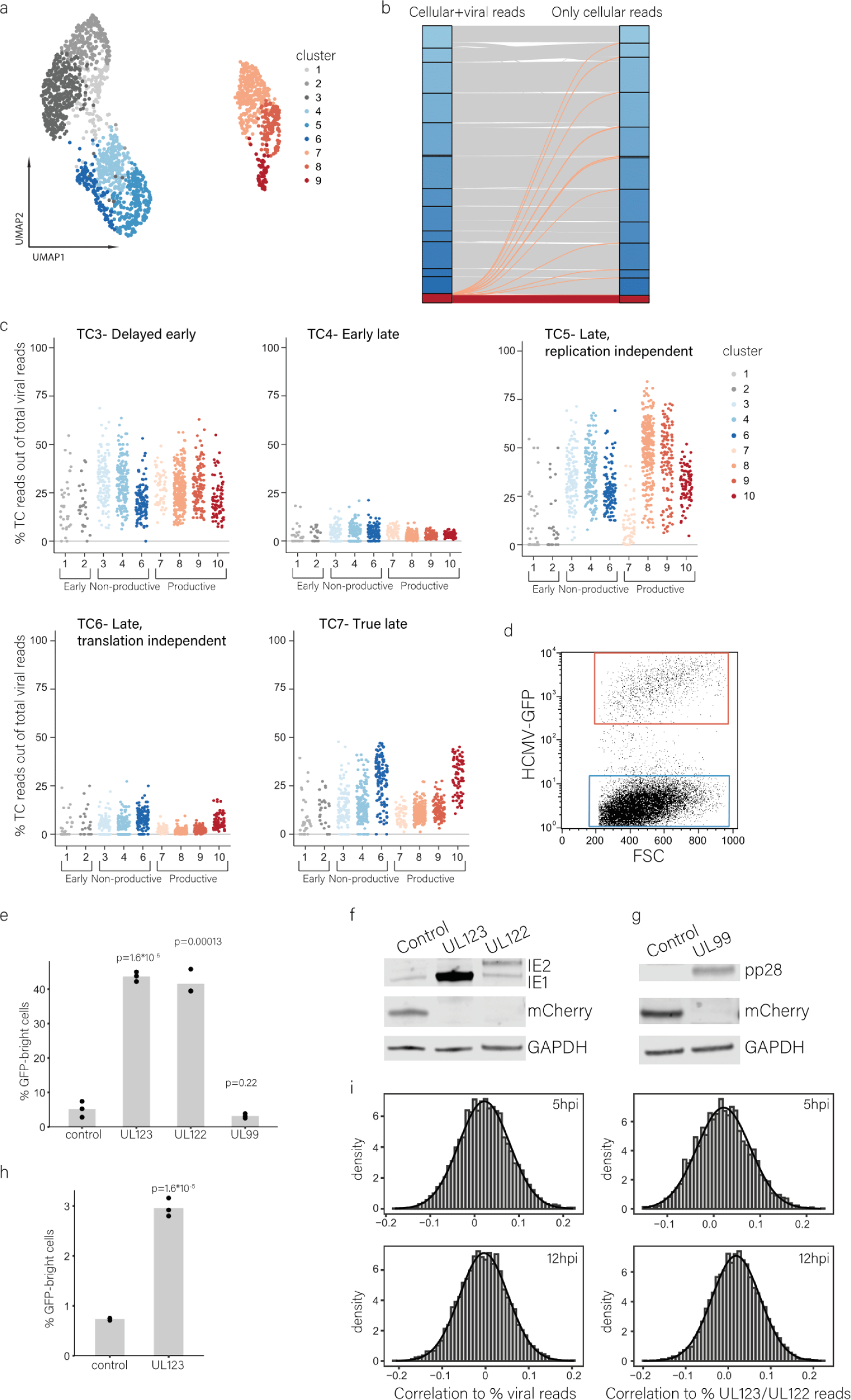
Expression of viral immediate early genes govern infection outcome of MDMs. **a**. Projection of HCMV-GFP infected MDMs at 5-144 hpi, analyzed based solely on cellular transcripts and colored by cluster assignment. **b**. Cluster assignment of each BAL myeloid cell according to analysis based on both viral and cellular genes (left) or on analysis based solely on cellular genes (right). Each cell is represented by a single line. The different non-productive cell clusters are colored in blue colors. Productively infected cells and clusters are colored in red. **c**. Relative expression levels, in the different MDM clusters, of viral genes from temporal classes 3- 7 based on ^28^. **d**. Flow cytometry analysis of THP1 derived macrophages (DMs) infected with HCMV-GFP. Analysis was performed at 3dpi. Blue and red gates mark the GFP-dim and GFP-bright populations, respectively. **e**. Percentage of productively infected cells in infected THP1 DMs ectopically expressing mCherry (control), UL123, UL122 or UL99. Points show measurement of biological replicates. p-values as calculated by t-test on the replicates are indicated. **f**. IE1/IE2 and mCherry protein levels at 8hpi visualized by western blot in THP1 DMs ectopically expressing UL123, UL122 or mCherry-flag as control. GAPDH is shown as loading control. **g**. pp28 (pUL99) and mCherry protein levels at 8hpi visualized by western blot in infected THP1 DMs ectopically expressing UL99 or mCherry-flag as control. GAPDH is shown as loading control. **h**. Percentage of productively infected cells in infected PMA-treated Kasumi-3 ectopically expressing UL123 or mCherry-flag as control. Points show measurement of biological replicates. p-values as calculated by t-test on the replicates are indicated. **i**. Distribution of the Spearman correlation coefficients between cellular transcript levels and viral gene expression level (left panels) or UL123 and UL122 expression levels (right panels), in early HCMV-GFP infected MDMs from 5hpi (top panels) or 12 hpi (bottom panels). The black line marks the matching normal distribution.

**Extended Data Figure 5.**
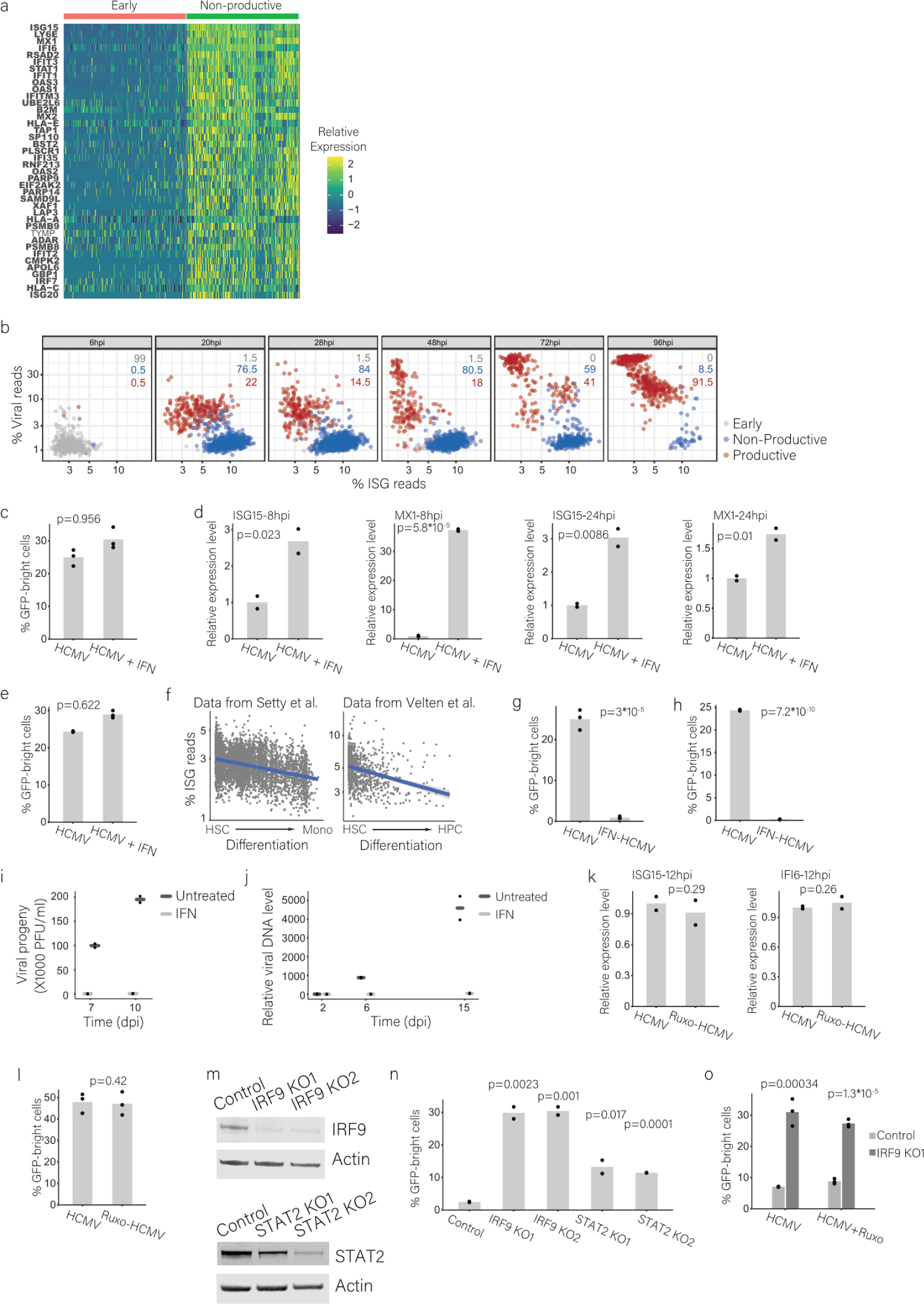
Expression of ISGs prior to infection but not their induction following infection affects infection outcome. **a**. Heatmap depicting the 40 most significantly increased genes in non-productive MDMs compared to early MDMs. ISGs are highlighted in bold. Z-scores of the log of the expression values are shown. **b**. Viral gene expression level versus ISG expression level in single HCMV- infected fibroblasts at different times post infection. Data taken from ^31^. The percentage of each group of cells in the given time point is indicated. **c.** Percentage of productively infected MDMs either untreated (HCMV), or treated with interferons (IFNs) at the time of infection (HCMV+IFN). Points show measurements of biological replicates. p-value of the decrease in productively infected MDMs following IFN treatment, as calculated by one tailed t-test on the replicates, is indicated. **d.** Relative expression level of representative ISGs, measured by qPCR, in infected MDMs either untreated (HCMV) or treated with IFNs at the time of infection (HCMV+IFN), at 8 and 24 hpi. Points show measurements of biological replicates. p-values as calculated by t-test on the replicates are indicated. **e.** Percentage of productively infected fibroblasts either untreated (HCMV), or treated with IFNs at the time of infection (HCMV+IFN). Points show measurements of biological replicates. p-value of the decrease in productively infected fibroblasts following IFN treatment, as calculated by one tailed t-test on the replicates, is indicated. **f.** ISG expression level versus differentiation status in single bone marrow human haematopoietic stem cells. Data taken from ^36^ shows differentiation trajectory from hematopoietic stem cells (HSCs) to monocytes (mono, left panel) and data from ^35^ shows differentiation trajectory from hematopoietic stem cells (HSCs) to hematopoietic progenitor cells (HPCs, right panel). The linear regression line with the confidence interval is shown. **g.** Percentage of productively infected MDMs either untreated (HCMV), or treated with IFNs 24 hours prior to infection (IFN-HCMV). Points show measurements of biological replicates. p-value of the decrease in productively infected MDMs following IFN treatment, as calculated by one tailed t-test on the replicates, is indicated. **h.** Percentage of productively infected fibroblasts either untreated (HCMV), or treated with IFNs 24 hours prior to infection (IFN-HCMV). Points show measurements of biological replicates. p-value of the decrease in productively infected fibroblasts following IFN treatment, as calculated by one tailed t-test on the replicates, is indicated. **i.** Measurements of infectious virus in supernatants from infected MDMs either untreated or treated with interferons (IFNs) 24 hours prior to infection, at different times post infection. **j**. qPCR measurements of viral genomes from infected MDMs either untreated or treated with interferons (IFNs) 24 hours prior to infection, at different times post infection. The plot shows the mean of 2 independent biological replicates. **k.** Relative expression level of representative ISGs, measured by qPCR, in infected MDMs either untreated (HCMV) or treated with ruxolitinib 24 hours prior to infection (Ruxo-HCMV), at 12 hpi. Points show measurements of biological replicates. p-values as calculated by t-test on the replicates are indicated. **l**. Percentage of productively infected MDMs either untreated (HCMV), or treated with ruxolitinib 24 hours prior to infection (Ruxo-HCMV). Points show measurements of biological replicates. p-value of the increase in productively infected MDMs following Ruxo treatment, as calculated by one tailed t-test on the replicates, is indicated. **m**. IRF9 or STAT2 levels as measured by western blot in THP1 DMs following CRISPR KO of IRF9 or STAT2, respectively. KO using two different gRNAs for each gene are shown. Actin was used as loading control. **n.** Percentage of productively infected THP1 DMs following CRISPR KO of a control gene, IRF9 or STAT2. Two independent gRNAs were used for IRF9 and STAT2. Points show measurements of biological replicates. p-value of IRF9 or STAT2 KO compared to the control, as calculated by t-test on the replicates is indicated. **o.** Percentage of productively infected THP1 DMs either untreated (HCMV) or treated with ruxolitinib at the time of infection (HCMV+Ruxo) following CRISPR KO of a control gene or IRF9. Points show measurements of biological replicates. p-value of IRF9 KO compared to the control, as calculated by t-test on the replicates, is indicated.

**Extended Data Figure 6.**
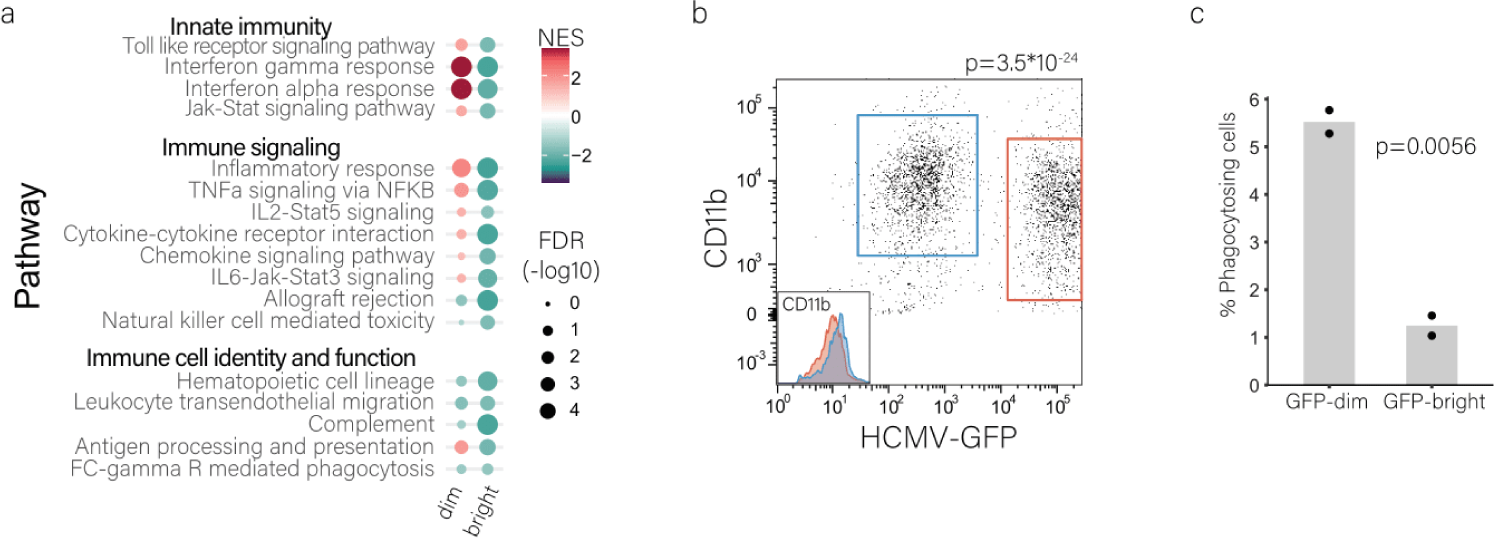
Effects of HCMV infection on macrophage cell identity. **a**. Gene set enrichment analysis (GSEA) of differentially expressed genes between GFP-dim and GFP-bright BAL infected macrophages compared to mock-infected cells, on innate immunity, immune signaling and immune identity and function pathways. **b.** Flow cytometry of CD11b surface levels in HCMV-GFP infected PMA-induced MDMs. The blue and red gates mark the non-productive and productive MDMs, respectively, as measured by GFP level. The inset in the bottom left shows the distribution of CD11b levels in the two populations. p value of the decrease in the mean fluorescence intensity in GFP-bright MDMs on the measured single cells is calculated using one-tailed t-test. Representative of two independent replicates is shown. **c.** Percentage of phagocytosing GFP-dim and GFP-bright HCMV-GFP infected M-CSF-induced MDMs. Points show measurements of biological replicates. p-value as calculated by t-test on the replicates is indicated.

**Extended Data Figure 7.**
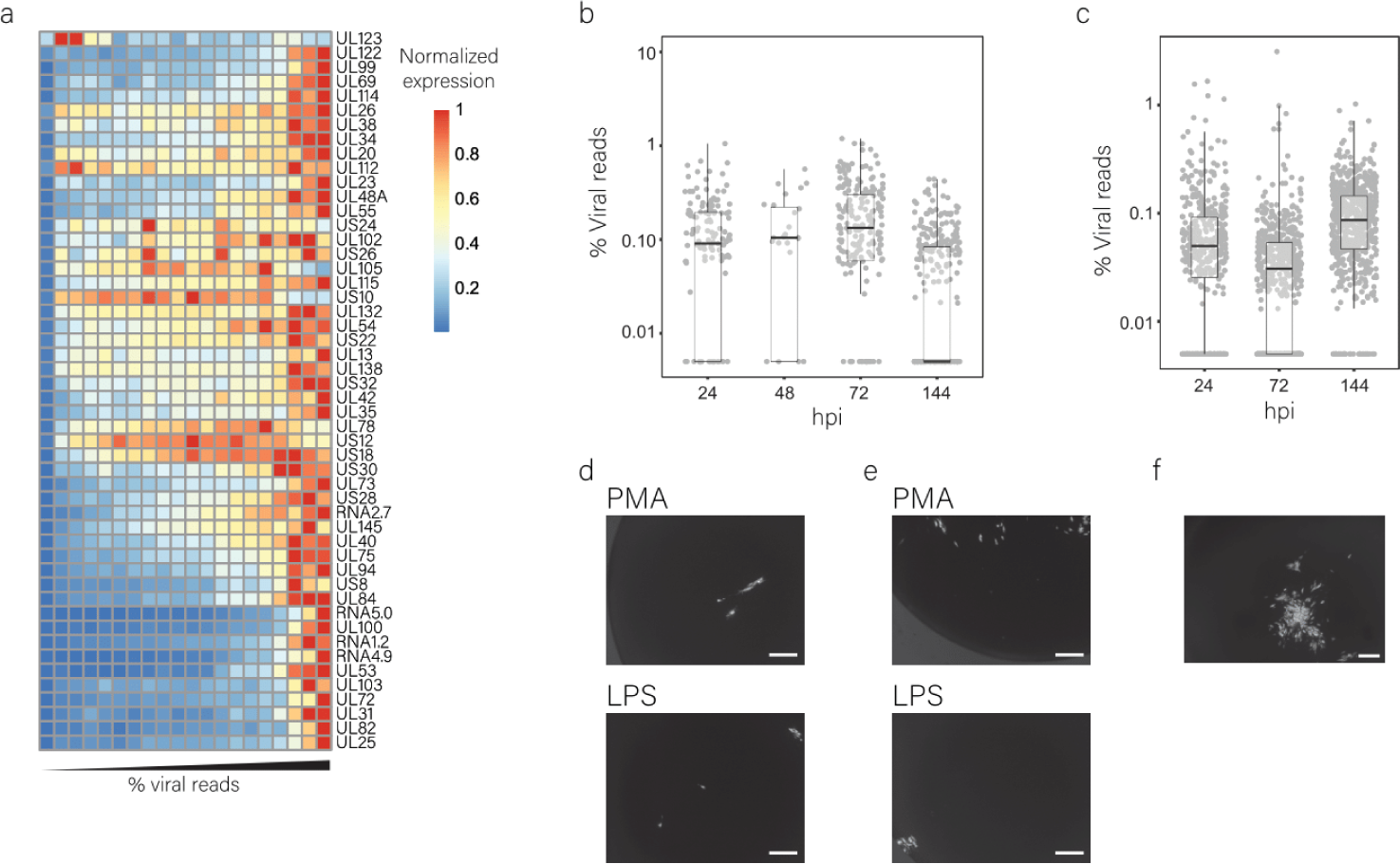
Non-productive MDMs continuously express viral transcripts and can reactivate. **a.** Heatmap depicting absolute levels of viral transcripts, in reads per million, in productive MDMs. Staging of the cells and ordering of the viral genes are as in Fig. 7a. The expression levels in the heatmap are relative to the highest level of each transcript. **b.** Percentage of viral reads in single monocytes along infection. **c.** Percentage of viral reads in single non-productive PMA-induced MDMs along infection. Data from 72 hpi and 144 hpi is from scRNA-seq of only GFP-dim cells to avoid noise coming from GFP-bright cells, expressing high levels of viral transcripts. **d.** Plaques formed on fibroblasts co-cultured with reactivated M-CSF-induced MDMs. Reactivation was done by either PMA or LPS treatment. Scale bars are 500μm. **e.** Plaques formed on fibroblasts co-cultured with reactivated PMA-induced MDMs. Reactivation was done by either PMA or LPS treatment. Scale bars are 500μm. **f.** Plaques formed on fibroblasts co-cultured with reactivated IFN pre-treated, PMA-induced MDMs. Reactivation was done by PMA. Scale bar is 500μm.

